# Neurotrauma-induced vulnerability of C9orf72 ALS iPSC-motor neurons revealed by biofidelic stretch injury

**DOI:** 10.1101/2024.03.21.586073

**Authors:** Eric J. Martin, Catrina Reyes, Angela Mitevska, Citlally Santacruz, Francesco Alessandrini, Ian E. Jones, Gopinath Krishnan, Gerard Esteruelas, Fen-Biao Gao, John D. Finan, Evangelos Kiskinis

## Abstract

A hexanucleotide repeat expansion (HRE) in *C9orf72* is the most common genetic cause of amyotrophic lateral sclerosis (ALS) and frontotemporal dementia (FTD). However, patients with the HRE exhibit a wide disparity in clinical presentation and age of symptom onset suggesting an interplay between genetic background and environmental stressors. Neurotrauma, resulting from traumatic brain or spinal cord injury has been shown to increase the risk of ALS in epidemiological studies. Here, we combine patient-specific induced pluripotent stem cells (iPSCs) with a custom-built device to deliver biofidelic stretch trauma to *C9orf72* patient and isogenic control motor neurons (MNs) *in vitro*. We find that mutant but not control MNs exhibit selective degeneration after a single incident of severe trauma, which can be partially rescued by pretreatment with a *C9orf72* antisense oligonucleotide. A single incident of mild trauma does not cause degeneration but leads to cytoplasmic accumulation of TDP-43 in *C9orf72* MNs. This mislocalization, which only occurs briefly in isogenic controls, is eventually restored in *C9orf72* MNs after 6 days. Lastly, repeated mild trauma ablates the ability of patient MNs to recover. These findings highlight alterations in TDP-43 dynamics in *C9orf72* ALS patient MNs following traumatic injury and demonstrate that neurotrauma compounds neuropathology in *C9orf72* ALS. More broadly, our work establishes an *in vitro* platform that can be used to interrogate the mechanistic interactions between ALS and neurotrauma.

## INTRODUCTION

Amyotrophic lateral sclerosis (ALS) is a progressive neurodegenerative disease with no effective therapeutic treatments [1]. It selectively affects upper and lower motor neurons (MNs), with cell death attributed to the disruption of an array of cellular processes including RNA metabolism, protein degradation pathways, cytoskeletal homeostasis, and neuronal excitability [2, 3]. Approximately 10% of ALS cases are caused by inherited genetic mutations, designated as familial ALS (fALS), while the remaining cases are classified as sporadic ALS (sALS) and likely arise from an interaction between genetic predisposition and environmental factors [4–6]. Roughly 30-50% of ALS patients also suffer from frontotemporal dementia (FTD), which is characterized by neurodegeneration in the frontal and temporal cortex and associated with speech and executive dysfunction [1]. The most prevalent genetic cause of ALS and FTD is an intronic hexanucleotide repeat expansion (HRE; GGGGCC_n_) in *C9orf72*, which is present in 40-50% of fALS and 5-10% of sALS cases, as well as up to 50% of all FTD cases [7–10]. The *C9orf72* HRE causes a reduction in C9orf72 protein levels and propagates deleterious gain-of-function effects, driven by the formation of C9-HRE RNA foci and the aggregation of dipeptide repeat (DPR) proteins that are transcribed and translated from the HRE likely through noncanonical mechanisms [11–17].

Patients carrying the *C9orf72* HRE are characterized by a diverse clinical presentation with a subset of individuals exhibiting only ALS or only FTD symptoms, while others are diagnosed with both [7–10]. The age of onset for C9orf72 patients is also highly variable and can range between 40 to 90 years, while in rare cases C9-HRE carriers never exhibit disease symptoms [18–23]. While the length of the C9-HRE can range between tens to thousands of repeats, the age of onset does not definitively correlate with any specific C9-HRE associated pathology such as repeat length, prevalence of RNA foci or DPR aggregates [15, 24–27]. Recent studies evaluating the penetrance of the C9orf72 repeat expansion suggest high variability in patient families ranging between 16 to 60% [28–30]. This pronounced heterogeneity suggests that genetic interactions and environmental determinants play a significant role in shaping disease prognosis and pathogenesis in C9-HRE ALS/FTD patients.

Traumatic brain injury (TBI) caused by a blow to the head or body has been repeatedly highlighted in epidemiological studies as a significant environmental risk factor for developing neurodegenerative diseases like ALS and FTD [31–33]. TBI is among the most prevalent neurological diagnoses, impacting as many as 3 million Americans annually [34, 35]. The clinical spectrum of TBI spans a wide continuum of severity that can include patients with enduring and chronic symptoms [36, 37]. Even instances of seemingly mild traumatic events have the potential to instigate downstream pathological changes in the brain and spinal cord. This has been demonstrated prominently in cases of chronic traumatic encephalopathy (CTE), a complex and progressive neurodegenerative disease caused by repetitive but usually mild brain injuries [38, 39]. The association between TBI and ALS/FTD is further exemplified by the shared neuropathology of TAR DNA-binding protein 43 (TDP-43) aggregation [40–43]. TDP-43 is a predominantly nuclear RNA binding protein involved in RNA transport and splicing, that becomes depleted from the nucleus and forms cytoplasmic aggregations in the overwhelming majority of ALS (∼97%) and as many as 50% of all FTD cases [44–46]. Notably, widespread TDP-43 aggregation is also seen in the brain and spinal cord of patients who have suffered TBI and CTE [47–49]. While these observations indicate a potential link between these diverse neurodegenerative diseases, it is unclear whether the mechanisms that drive TDP-43 pathology in ALS and TBI are shared or divergent. What also remains unclear is whether neurotrauma resulting from traumatic brain or spinal cord injury can trigger ALS disease pathophysiology and conversely, whether an ALS genetic background confers heightened vulnerability to neurotrauma, specifically in the context of human neurons.

To address these critical gaps in knowledge we established an *in vitro* model system of *C9orf72* ALS patient-specific induced pluripotent stem cell (iPSC) derived motor neurons (MNs), which can be physically traumatized by applying highly controlled and biofidelic stretch-induced injury [50–52]. We find that the *C9orf72* HRE renders patient-derived MNs more susceptible to mild and severe traumatic injury reflected in selective degeneration and altered TDP-43 localization dynamics. These pathologies, which were considerably milder in isogenic control MNs, could be partially rescued by preemptive administration of an antisense oligonucleotide (ASO) targeting the C9-HRE. We also find that repeated mild trauma ablates the ability of C9-HRE patient MNs to effectively recover and causes TDP-43 pathology and degeneration. These findings highlight alterations in TDP-43 dynamics in *C9orf72* ALS patient MNs following traumatic injury and establish an *in vitro* platform that can be used to interrogate the mechanistic interactions between ALS and neurotrauma.

## RESULTS

### Establishing an *in vitro* model of neurotrauma

The physical injury of the brain or the spinal cord can cause primary localized defects including neurodegeneration, axonal severing and edema followed by a range of downstream pathological conditions such as excitotoxicity and inflammation [34, 35]. Diffuse axonal injury (DAI) that ranges between stretching and shearing is caused by the impact of the mechanical force and the vigorous shaking of the brain and the spinal cord within the central nervous system [53–57]. To simulate neurotrauma and specifically DAI *in vitro,* we used a recently developed instrument that can deliver biofidelic stretch injury to cultured neurons [52, 58].

We selected a well-characterized *C9orf72* ALS/FTD patient iPSC line and its corresponding isogenic control (Cell line CS29; Table 1 and Methods), generated by CRISPR/Cas9 gene editing, and differentiated them into ISL1/2/MAP2 positive MNs using established protocols (Fig. 1A, Supplementary Fig. 1A-C) [59]. Upon differentiation, MNs were plated into 96-well plates with a flexible polydimethylsiloxane (PDMS) bottom and a feeder layer of primary mouse glial cells. To traumatize the cultures of MNs, plates were positioned above a post array that allowed high-speed, regulated trauma at controlled displacements of individual wells (Fig. 1B-C) [52, 58]. Upward drive of the post array caused extension of the flexible PDMS membrane causing it to stretch in the horizontal plane (Supplemental Video S1). Critically, stretch trauma of one well does not affect adjacent wells, allowing direct side-by-side comparisons of conditions (Supplemental Video S2), while the instrument can inflict injury of varying degrees. A detailed experimental characterization of this neurotrauma model is based on measuring nominal indentation depth values (mm displacement) and resulting membrane strains (Lagrangian strain) [58]. The quantification of this interaction showed that an indentation depth of 1 mm, induced a 9.6 ± 0.7% strain, while a 3 mm displacement induced a 66.9 ±4.4% strain (Supplementary Fig. 1D). These values are consistent with the best available estimates of brain tissue strains in mild and severe injury respectively [60–62]. A recent study that reconstructed impact biomechanics in collisions that led to concussion in football players (i.e. mild injuries) found that a strain value of 10% was the threshold that best predicted injury [63]. A study that reconstructed more severe injuries occurring in car crashes using human and non-human primate data showed a 25% strain to be the tolerance associated with severe injury [64]. Thus, based on published findings [52, 58, 65] and our calculations of Lagrangian strain in our model, we classified displacements of 1.5 mm and ≥ 3 mm as mild and severe, respectively (Fig. 1D). We also quantified the consistency of the membrane strain across repeated indentations to assess the feasibility of a repeated injury protocol. The membrane strain did not decline in a statistically significant manner over the course of 10 consecutive 2 mm deep indentations (Supplementary Fig. 1E).

**Table 1:**

| Line | Source | Disease | Method of Correction | Sex | Age | Link |
| --- | --- | --- | --- | --- | --- | --- |
| CS29<br>*C9orf72*<br>ALS | Cedars-Sinai<br>Biomanufacturing<br>Center | *C9orf72*<br>ALS |  | M | 47 | CS29 ALS |
| CS29<br>Isogenic<br>Control | Cedars-Sinai<br>Biomanufacturing<br>Center | Non-<br>disease<br>control | CRISPR/Cas9 | M | 47 | CS29 ISO |
| CS52<br>*C9orf72*<br>ALS | Cedars-Sinai<br>Biomanufacturing<br>Center | *C9orf72*<br>ALS |  | M | 49 | CS52 ALS |
| CS52<br>Isogenic<br>Control | Cedars-Sinai<br>Biomanufacturing<br>Center | Non-<br>disease<br>control | CRISPR/Cas9 | M | 49 | CS52 ISO |

**Figure 1.**
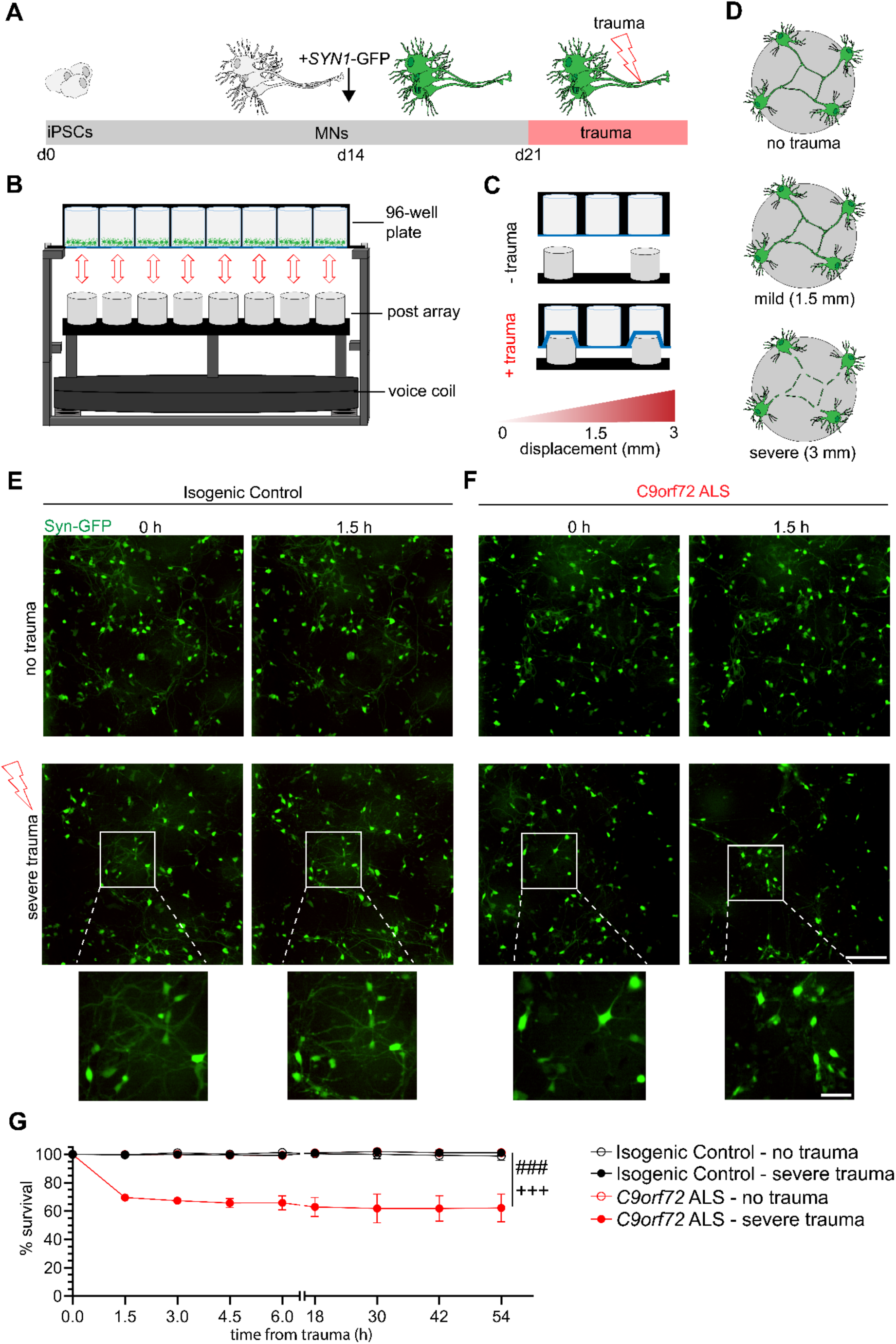
*C9orf72* ALS motor neurons show selective degeneration following severe stretch trauma. **(A)** Differentiation schematic. Motor neurons are generated from iPSCs across 14 days, then transduced with *SYN1*-GFP. One week after transduction, trauma was performed. **(B)** Experimental apparatus for induction of stretch trauma. A voice coil drives the post array into a PDMS-bottom plate. **(C)** Visual schematic of stretch trauma. PDMS stretches as posts push up before returning to original conformation. **(D)** Visual schematic of varying degrees of trauma. **(E-F)** Live imaging of CS29 *SYN1*-GFP motor neurons immediately prior (0 h) and 1.5 hours after either no trauma control (NTC) or severe trauma (3 mm) for either isogenic control (**E**) or *C9orf72* (**F**) motor neurons. Cutout shows neuralization after severe trauma. Scale bar = 200 μm; inset scale bar = 50 μm **(G)** Quantification of percent survival of neurons within the frame of imaging across intervals of 1.5 hours for the first 6 hours and every 12 hours thereafter for a total of 54 hours. Experimental averages of 3 independent differentiations with 245-350 neurons from 8 total fields of view. *** = p < .0005 for two-way ANOVA. Tukey’s multiple comparison tests for *C9orf72* ALS – 3mm: p < 0.005 compared to Isogenic Control – no trauma (###) and Isogenic Control – 3mm (+++). Data are represented as mean ± SEM.

### *C9orf72* ALS MNs selectively degenerate following severe stretch trauma

To specifically visualize iPSC-derived MNs using live imaging analysis, we transduced the cultures with a lentivirus reporting GFP under the Synapsin 1 promoter (*SYN1*-GFP) following their plating on PDMS and prior to injury (Fig. 1A). We first asked whether a single incident of severe trauma (3 mm) would inflict differential responses between patient and isogenic control MNs. Quantification of *SYN1*-GFP cells revealed that *C9orf72* ALS MNs displayed a significant 30% reduction in cell number at 1.5 hours post-trauma relative to the no trauma control condition, followed by a persistent gradual decline reaching 40% approximately 2 days later (no trauma control *Vs*. severe trauma in *C9orf72*; N=3, p<0.0001) (Fig. 1E-G). In contrast, the isogenic control MNs injured at the same time exhibited no significant changes even after 2 days (no trauma control *Vs*. severe trauma in controls; N=3, p=NS) (Fig. 1E-G). To determine if the higher vulnerability of mutant C9orf72 MNs to stretch trauma was reproducible across other cell lines we performed the same experiment using a second mutant *C9orf72* iPSC line and its corresponding isogenic control (Cell line CS52; Table 1 and Supplementary Fig. 2). Although we did not observe significant effects at 3mm displacement, both genotypes were sensitive to a more severe stretch injury of 3.6mm, with the mutant C9orf72 MNs exhibiting a significantly more pronounced reduction in MN numbers relative to their isogenic controls (Supplementary Figure 2). These data demonstrate that the ALS-associated C9-HRE renders MNs susceptible to severe stretch trauma.

To ascertain whether the heightened susceptibility to severe trauma was intrinsically linked to the *C9orf72* HRE, we subjected the MN cultures to a pre-treatment with either a scrambled or an ASO targeting the sense *C9orf72* repeat expansion and repeated the experimental paradigm of severe injury (Fig. 2A-B). Consistent with previous reports [66], we found that the ASO reduced the expression of both the sense repeat and total *C9orf72* transcript by approximately 70% and 30% respectively, relative to samples treated with a scrambled, control ASO (Fig. 2C). Monitoring of *SYN1*-GFP neurons 1.5 hours after trauma revealed a 30% reduction in the *C9orf72* cultures, whereas as previously observed the isogenic controls were unaffected (control trauma *Vs*. C9 trauma; N=3, p< 0.0001) (Fig. 2B, 2D). Notably, pretreating the *C9orf72* ALS MNs with the HRE-targeting ASO caused a significant but incomplete rescue of degeneration following severe trauma, suggesting that the sense repeat RNA is in part responsible for MN vulnerability to neurotrauma (scrambled ASO trauma *Vs*. C9 ASO trauma in mutant *C9orf72* MNs; N=3, p< 0.005) (Fig. 2B, 2D).

**Figure 2.**
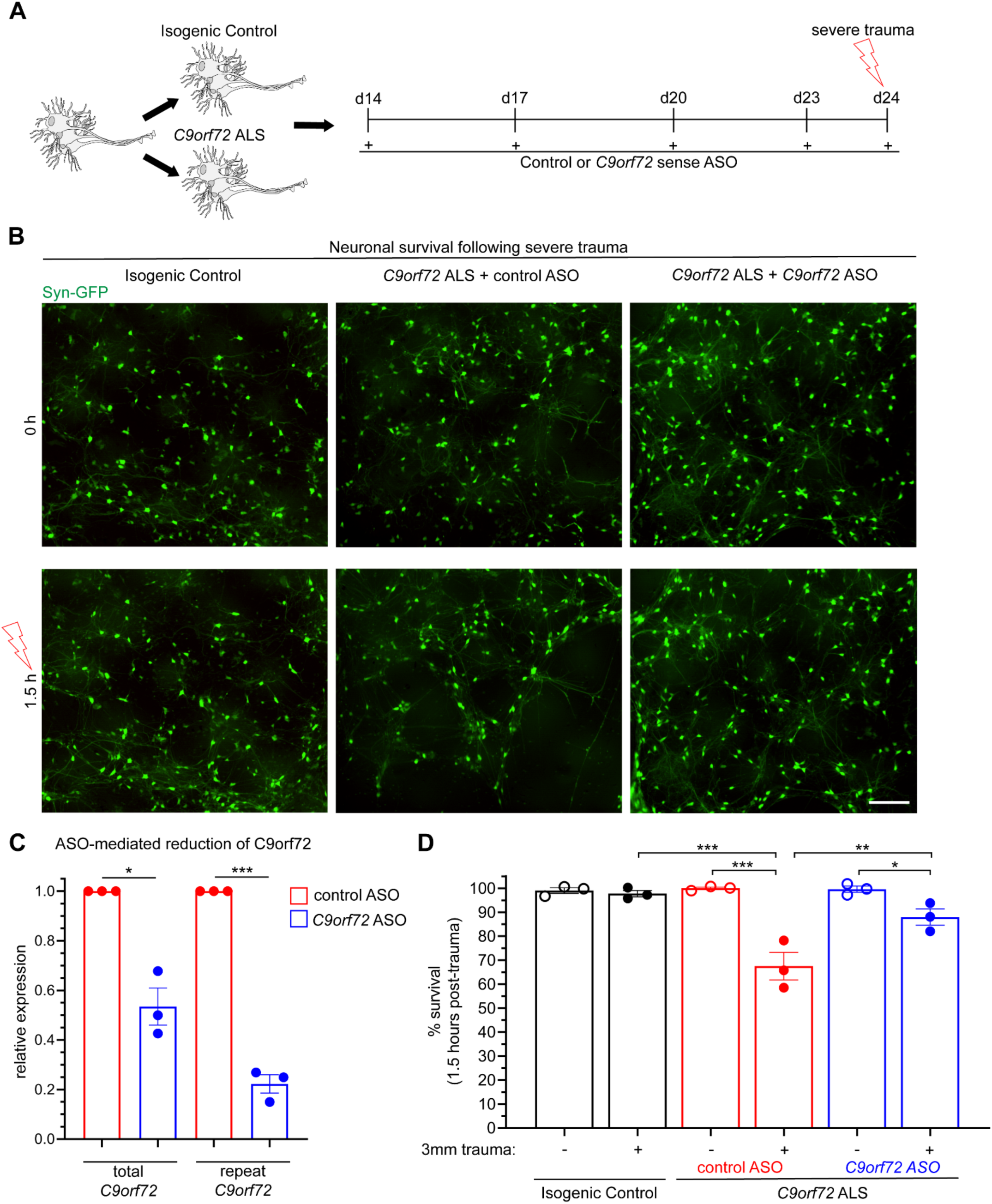
Neurodegeneration following severe trauma is partially rescued by pretreatment with an ASO targeting the C9orf72 repeat expansion. **(A)** Experimental schematic for ASO treatment prior to severe trauma. **(B)** Live imaging of CS29 *SYN1*-GFP MNs 1.5 hours following severe trauma or no trauma. Scale bar = 200 μm. **(C)** Relative expression of total *C9orf72* and repeat-associated *C9orf72* in *C9orf72* ALS MNs following treatment from a *C9orf72* ASO specifically targeting the repeat expansion compared to a scramble ASO from 3 biological replicates. **(D)** Quantification of percent survival 1.5 hours following trauma, showing differences between *C9orf72* ALS pretreated with a scramble ASO (red) and a repeat-targeting ASO (blue). Experimental averages of 3 independent differentiations from 16 fields of view quantifying 237-363 total neurons. Two-way ANOVA p < 0.0005. For Tukey’s multiple comparisons test, * = p < 0.05 ** = p < 0.01 *** = p < 0.005. Data are represented as mean ± SEM.

### Mild stretch trauma increases the abundance of G_4_C_2_ RNA foci but not GP-DRP in *C9orf72* ALS MNs

Having established an increased susceptibility of *C9orf72* MNs to severe trauma, we next sought to characterize pathological changes under conditions of trauma that did not elicit immediate degeneration. We first examined *C9orf72* and isogenic control MN cultures over an extended time course of over 144 hours (6 days) after a single incidence of mild traumatic injury (1.5 mm). This analysis showed that neither mutant nor control MNs were affected, indicating the absence of a delayed or secondary wave of degeneration (no trauma *Vs*. mild trauma; N=3, p=NS) (Fig. 3A-C). We next interrogated the effects of mild sublethal injury to *C9orf72*-associated gain-of-function neuropathology by conducting FISH and ELISA assays targeting C9-HRE RNA foci and poly-GP DPR production, respectively (Fig. 3D). As expected, the abundance of nuclear (G_4_C_2_)n RNA foci was found to be significantly higher in C9-HRE patient MNs at baseline. Mild trauma induced a 50% increase in the number of C9-HRE patient MNs with G_4_C_2_ RNA foci after 4 days, while the controls were unaffected (no trauma *Vs*. mild trauma in *C9orf72*; N=3, p< 0.0001) (Fig. 3E-F and Supplementary Fig. 3). Notably, pretreating the C9-HRE patient MNs with the sense targeting C9orf72 ASO prevented the stretch trauma-induced accumulation of HRE-RNA foci (control ASO *Vs*. C9-HRE *ASO*; N=3, p< 0.005) (Supplementary Fig. 3C). Intriguingly, the stretch-induced increase in RNA foci was not accompanied by a corresponding increase in the relative amount of poly-GP, suggesting that trauma does not induce non-ATG dependent translation unlike other forms of cellular stress (no trauma *Vs*. mild trauma in *C9orf72*; N=3, p=NS) (Fig. 3G) [67].

**Figure 3.**
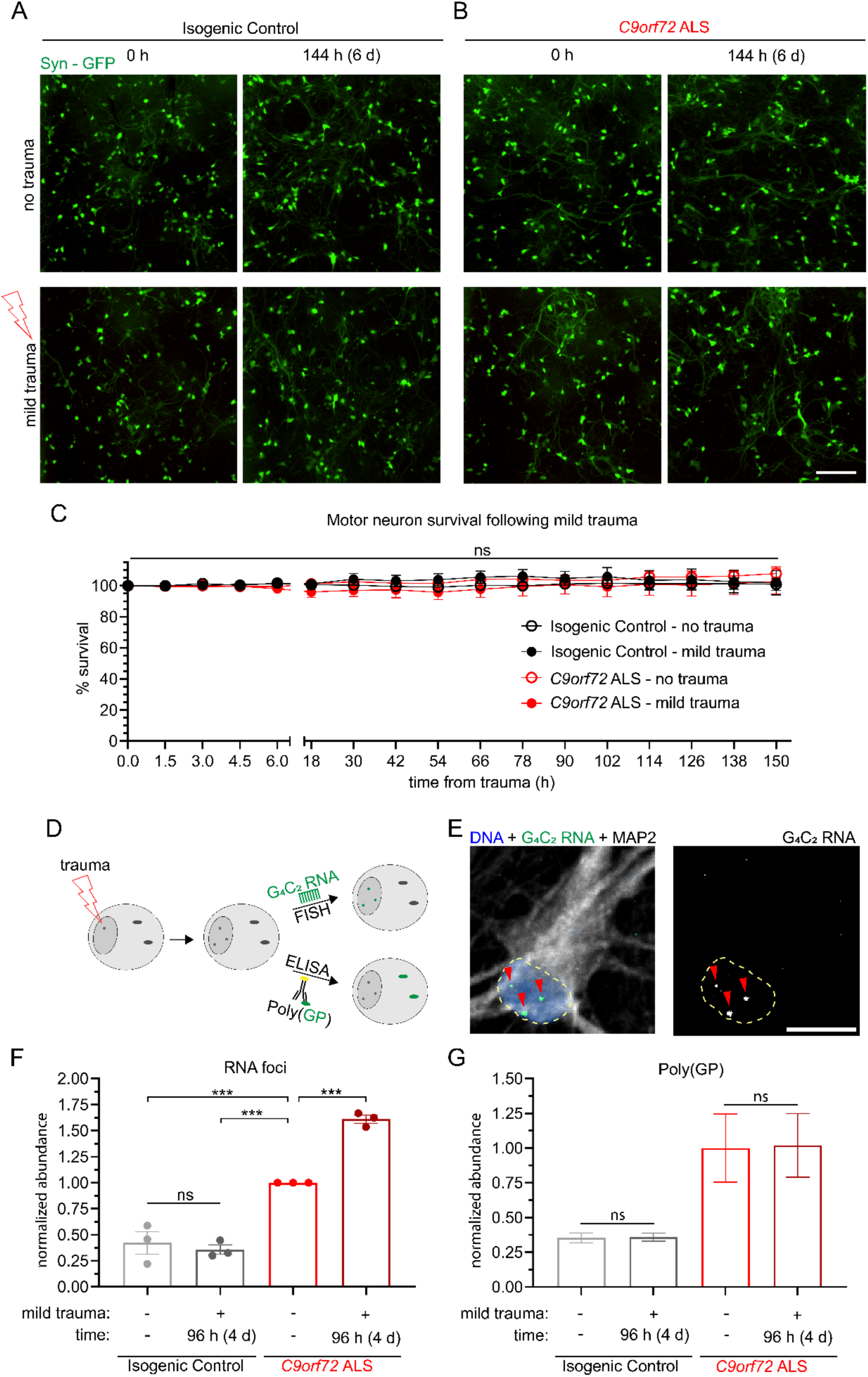
Mild trauma increases incidence of RNA foci but not DPR production in C9orf72 ALS motor neurons. (A-B) Live imaging of CS29 synapsin-GFP isogenic control motor neurons across a period beyond 6 days (150 hours) following either no trauma or mild trauma (1.5 mm) in either isogenic control (**A**) or *C9orf72* ALS (**B**) motor neurons. Scale bar = 200 μm. **(C)** Quantification of percent survival for over 6 days (150h) following mild trauma. Two-way ANOVA NS. **(D)** Experimental schematic for quantification of toxic gain of function *C9orf72* pathologies. **(E)** Representative image of FISH probe analyzed for RNA foci. **(F)** Quantification of percent of neurons with RNA foci 4 days following mild trauma relative to *C9orf72* ALS NTC. Experimental averages for 3 independent differentiations from 147-205 neurons. Two-way ANOVA p < 0.0005. Tukey’s multiple comparisons test *** = p < 0.005 relative to *C9orf72* ALS NTC. **(G)** ELISA quantification of poly(GP) DPR 4 days following mild trauma. Data are represented as mean ± S

### Stretch trauma induces alterations in the nucleocytoplasmic localization of TDP-43

The cytoplasmic mis-localization of TDP-43 constitutes an essential convergent pathology between ALS and neurotrauma resulting from TBI, as observed in CTE cases [68, 69]. To assess the dynamics of TDP-43 in the context of the *in vitro* stretch injury model, we next subjected C9-HRE patient and isogenic control MNs to trauma spanning a spectrum of severities (0, 1, 1.5, 2, 2.5, and 3 mm membrane displacement) and quantified the nucleocytoplasmic (N/C) localization of TDP-43 by immunocytochemistry and confocal imaging 4 hours post-trauma (Fig. 4A, Supplementary Fig. 4A-B). Importantly, we did not observe any significant differences in the N/C localization of TDP-43 in the absence of trauma between mutant C9orf72 ALS and control MNs (C9orf72 *Vs*. control; N=3, M=72-84 cells, p=NS) (Supplementary Fig. 4C). To account for the variation in raw TDP-43 localization values across independent differentiations and trauma severities we used z-score standardization for all subsequent analyses (see Methods). The control MNs did not exhibit any significant alteration in the N/C ratio of TDP-43 across any of the trauma severities we examined (no trauma *Vs*. trauma in controls; N=3, M=84-91 cells, p=NS) (Fig. 4B-C and Supplementary Fig. 4A and D). In contrast, we found a significant reduction in the N/C ratio of TDP-43 in response to trauma equal or stronger to 1.5 mm in C9-HRE patient-derived MAP2 positive MNs (no trauma *Vs*. trauma in *C9orf72*; N=3, M=58-88 cells, p<0.005) (Fig. 4D-E and Supplementary Fig. 4B and E). Notably, this shift in N/C TDP-43 signal was not dose-dependent but rather threshold dependent.

**Figure 4.**
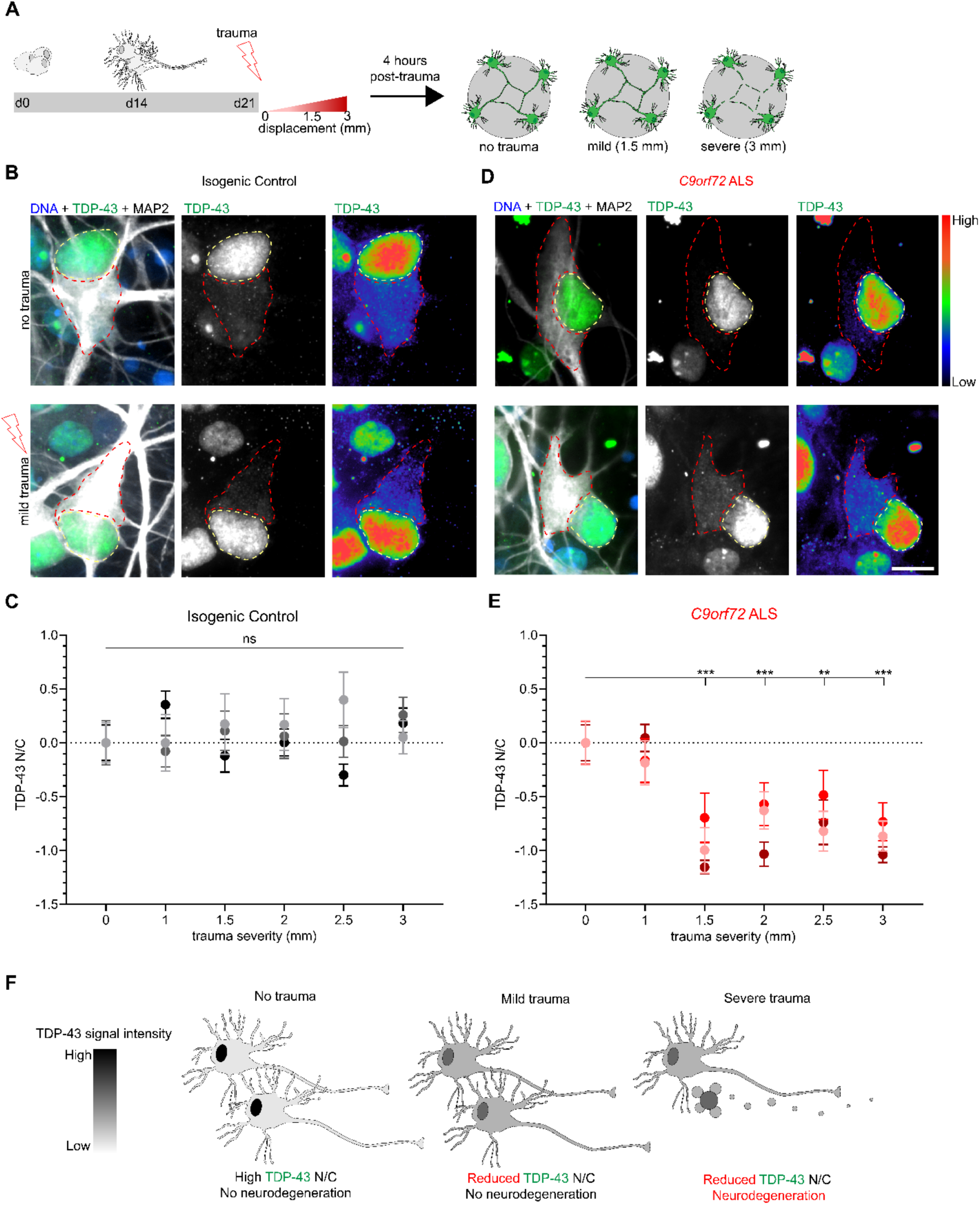
Traumatic injury induces TDP-43 mislocalization across a range of severities in C9orf72 ALS but not isogenic control motor neurons. **(A)** Schematic depicting paradigm for stretch trauma across a range of severities. **(B and D)** Immunocytochemistry shows TDP-43 distribution in either CS29 isogenic control (**B**) or *C9orf72* ALS **(D)** motor neurons prior to and 4 hours post-trauma. No trauma compared to 1.5mm displacement. Scale bar = 10 μm. **(C and E)** Z-score quantification of TDP-43 N/C at each displacement value relative to no trauma in either isogenic control (**C**) or *C9orf72* ALS (**E**) motor neurons. Data represent experimental averages from 3 independent differentiations for a total of 74 - 91 (**C**) or 58 – 87 (**E**) cells. (**C**) One-way ANOVA NS (**E**) One-way ANOVA p < 0.0005. Tukey’s multiple comparisons test *** p < 0.005, ** p < 0.01. Data are represented as mean ± SEM. **(F)** Schematic representation of mild and severe trauma compared to no trauma with respect to TDP-43 N/C and viability.

To further assess the specificity of this effect, we interrogated mutant *SOD1^A4V/+^* and isogenic control MN cultures derived from a well characterized set of ALS patient iPSC lines [70–72] that were subjected to the same wide range of traumatic injury paradigm (Supplementary Fig. 5). Crucially, the *SOD1^A4V/+^* MNs did not display any discernable reduction in the N/C TDP-43 ratio in these experiments (no trauma *Vs*. trauma in *SOD1^A4V/+^*; N=3, M=91-101 cells, p=NS), in accordance with the fact that *SOD1* ALS patients fall within the 3% of all patients that do not develop neuropathological TDP-43 aggregates postmortem [73, 74].

Our findings demonstrate that TDP-43 exhibits a significant shift in N/C localization 4 hours after severe or even mild traumatic injury without any measurable effects on neuronal viability, but only in the context of a relevant ALS genetic background (Fig. 4F). To better define the dynamics of this mis-localization we quantified the N/C ratio of TDP-43 in C9-HRE patient and isogenic control MNs with higher temporal resolution after a single mild traumatic insult. We found that isogenic control MNs exhibited a brief reduction in TDP-43 N/C ratio 15 minutes post-trauma that normalized to baseline levels by 30 minutes (no trauma *Vs*. trauma in controls at 15 min; N=3, M=177 cells, p<0.05) (Fig. 5A-B, Supplementary Fig. 6). In contrast, this post-traumatic shift in TDP-43 was delayed in *C9orf72* ALS MNs and did not occur until 4 hours (no trauma *Vs*. trauma in *C9orf72* at 4 hrs; N=3, M=170 cells, p<0.01) (Fig. 5C-D) as we previously described (Fig. 4D). The observation of a brief shift in TDP-43 localization followed by a full recovery in control MNs prompted us to next ask whether a similar recovery occurs at all in mutant *C9orf72* ALS MNs. Thus, we repeated the mild traumatic injury and quantified the localization of TDP-43 for up to 6 days post injury. The control MNs exhibited a consistent N/C ratio in TDP-43 throughout this extended time course (Fig. 5E-F). The *C9orf72* ALS MNs showed a sustained level of TDP-43 mis-localization, that peaked at 4 hours and recovered to pre-traumatic levels only after 6 days (no trauma *Vs*. trauma in *C9orf72* at 4 hrs p<0.01; at 96 hours p<0.01; at 144 hours p=NS; N=3, M=127,133, and 124 cells respectively) (Fig. 5G-H and Supplementary Fig. 7).

**Figure 5.**
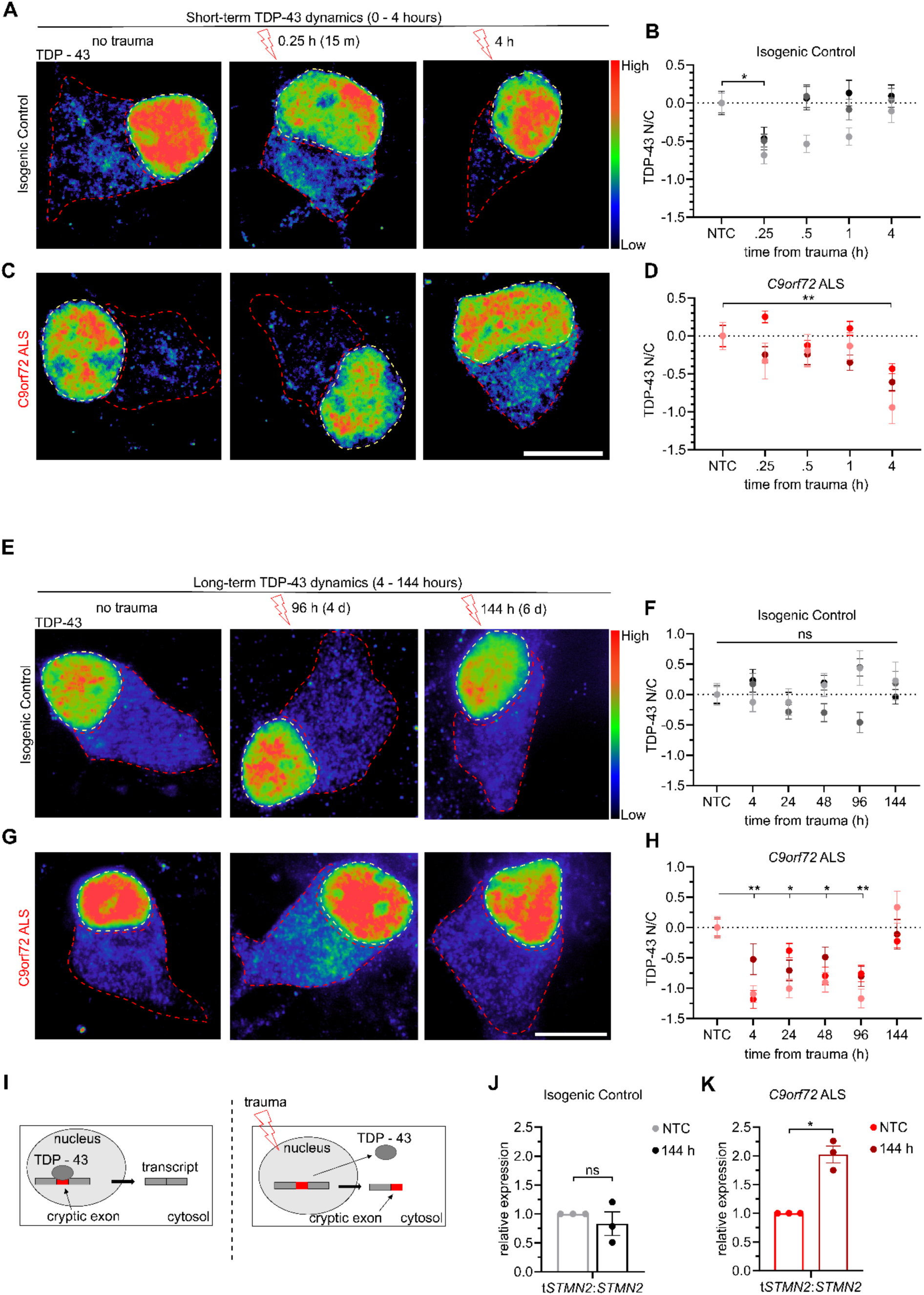
Mild trauma induces extended TDP-43 mislocalization and increased mis-splicing of STMN2 in *C9orf72* ALS motor neurons. (A and C) Immunocytochemistry showing TDP-43 in either CS29 isogenic control (**A**) or *C9orf72* ALS (**C**) motor neurons at 0 hours, 15 minutes (0.25 h), and 4 hours following mild trauma. Scale bar = 10 μm **(B and D)** Z-score quantification of TDP-43 N/C relative to no trauma in either isogenic control (**B**) or *C9orf72* ALS (**D**) motor neurons. Data represent experimental averages from 3 independent differentiations for a total of 133 - 162 (**B**) or 132 - 139 (**D**) cells. **(B)** One-way ANOVA p < 0.05. Tukey’s multiple comparisons test * = p < 0.05. (**D**) One-way ANOVA p < 0.05. Tukey’s multiple comparisons test ** = p < 0.01. **(E and G)** Immunocytochemistry showing TDP-43 in either isogenic control (**E**) or *C9orf72* ALS (**G**) motor neurons at 0 hours, 96 hours, and 144 hours following mild trauma. Scale bar = 10 μm **(F and H)** Z-score quantification of TDP-43 N/C relative to NTC in either isogenic control (**F**) or *C9orf72* ALS (**H**) motor neurons. Data represent experimental averages from 3 independent differentiations for a total of 118 – 133 (**F**) or 123 - 138 (**H**) cells. (**F**) One-way ANOVA NS. (**H**) One-way ANOVA p < .0005. Tukey’s multiple comparisons test * = p < 0.05, ** = p < 0.01. **(I)** Visual schematic of mis-splicing caused by mild trauma. **(J and K)** Quantification of relative expression of truncated *STMN2* (t*STMN2*) relative to total expression (*tSTMN2/STMN2*) by qPCR for Isogenic Control (**J**) or *C9orf72* ALS (**K**). Data for each line are normalized relative to no trauma. Data represent 3 independent differentiations analyzed by Welch’s t-test. * = p < 0.05. Data are represented as mean ± SEM.

Dysfunction of TDP-43 has recently been shown to increase the mis-splicing of target mRNAs that appear to be regulated post-transcriptionally such as *STMN2*, which encodes a regulator of microtubule homeostasis, [75–77]. Binding of TDP-43 to these mRNAs prevents mRNA decay and downregulation (Fig. 5I) To determine whether the prolonged mis-localization of TDP-43 induced by mild traumatic injury recapitulated aberrant RNA splicing, we employed qRT-PCR to quantify the level of cryptic exon inclusion in *STMN2* [77, 78]. We found that in the isogenic control MN cultures the relative proportion of truncated *STMN2* (*tSTMN2*) relative to full-length *STMN2* was unchanged 144 hours after injury (Fig. 5J). In contrast, *tSTMN2/STMN2* was moderately but significantly increased 144 hours following trauma in *C9orf72* ALS MNs (Fig. 5K). Notably, this occurs even though the N/C distribution of TDP-43 has returned to baseline levels at this point, suggesting that neurotrauma can cause lingering defects in mRNA splicing even beyond recovery of nuclear localization of TDP-43. Taken together, these findings underscore the potential for TDP-43 mis-localization to be a normative response to traumatic injury, as it also occurs in healthy control MNs, albeit briefly, and without evidence of truncated *STMN2*. However, the significantly delayed dynamics of the response and ultimate delayed recovery and sustained dysfunction of TDP-43 in C9-HRE MNs indicates that the ALS genetic background compromises the response to neurotrauma.

### Repetitive mild stretch trauma causes neurodegeneration and prevents recovery of TDP-43 mis-localization in *C9orf72* ALS MNs

The resilience of control MNs to stretch induced injury as well as the ability of *C9orf72* ALS MNs to ultimately restore TDP-43 localization following mild injury prompted us to investigate their susceptibility and molecular responses to repeated incidences of trauma. We designed an experimental paradigm where MNs were injured mildly on day 21, analyzed and allowed to recover for 6 days, injured again using the same mild force (1.5 mm), followed by another 6 days of recovery and analysis (Fig. 6A). MNs were afforded a 6-day recovery period to match the duration for TDP-43 recovery following acute trauma, while downstream pathologies like C9-HRE RNA foci and aberrant mRNA splicing may persist (Figs. 3F and 5K). Intriguingly, introducing a second instance of mild trauma proved sufficient to significantly reduce survival of *C9orf72* ALS MNs by approximately 10% (no trauma control *Vs*. 2x mild trauma in C9orf72; N=3, p<0.05), while isogenic control MNs continued to be resilient, as their numbers remained unaffected (no trauma control *Vs*. 2x mild trauma in controls; N=3, p=NS) (Fig. 6B-E). Quantification of the N/C localization of TDP-43 in control MNs showed no effect after a single or a second incidence of mild trauma (no trauma control *Vs*. 1x or 2x mild trauma in controls; N=3, M=204 and 211 cells, p=NS) (Fig. 7A, C and Supplementary Fig. 8). In accordance with this observation, there was no shift in the *tSTNM2/STMN2* transcript ratio (Fig. 7D). At the same time, in *C9orf72* ALS MNs the N/C distribution of TDP-43 was restored 6 days after one mild hit (Fig. 7B, E) as we observed previously (Fig. 5G-H). However, a subsequent mild hit compromised their ability to effectively recover and caused substantial reduction in N/C TDP-43 signal (no trauma control *Vs*. 1x mild trauma in *C9orf72* p=NS; Vs. 2x p<0.05; N=3, M=196 cells) (Fig. 7B, E and Supplementary Fig. 8). Critically, this defect was reflected in the abundance of *tSTMN2*, with a significantly increased accumulation after a second mild traumatic injury relative to a single acute mild trauma (Fig. 7F). Collectively, these data highlight the compounding effect of multiple injuries of a mild nature on neurodegenerative molecular pathologies and susceptibility of mutant *C9orf72* MNs.

**Figure 6.**
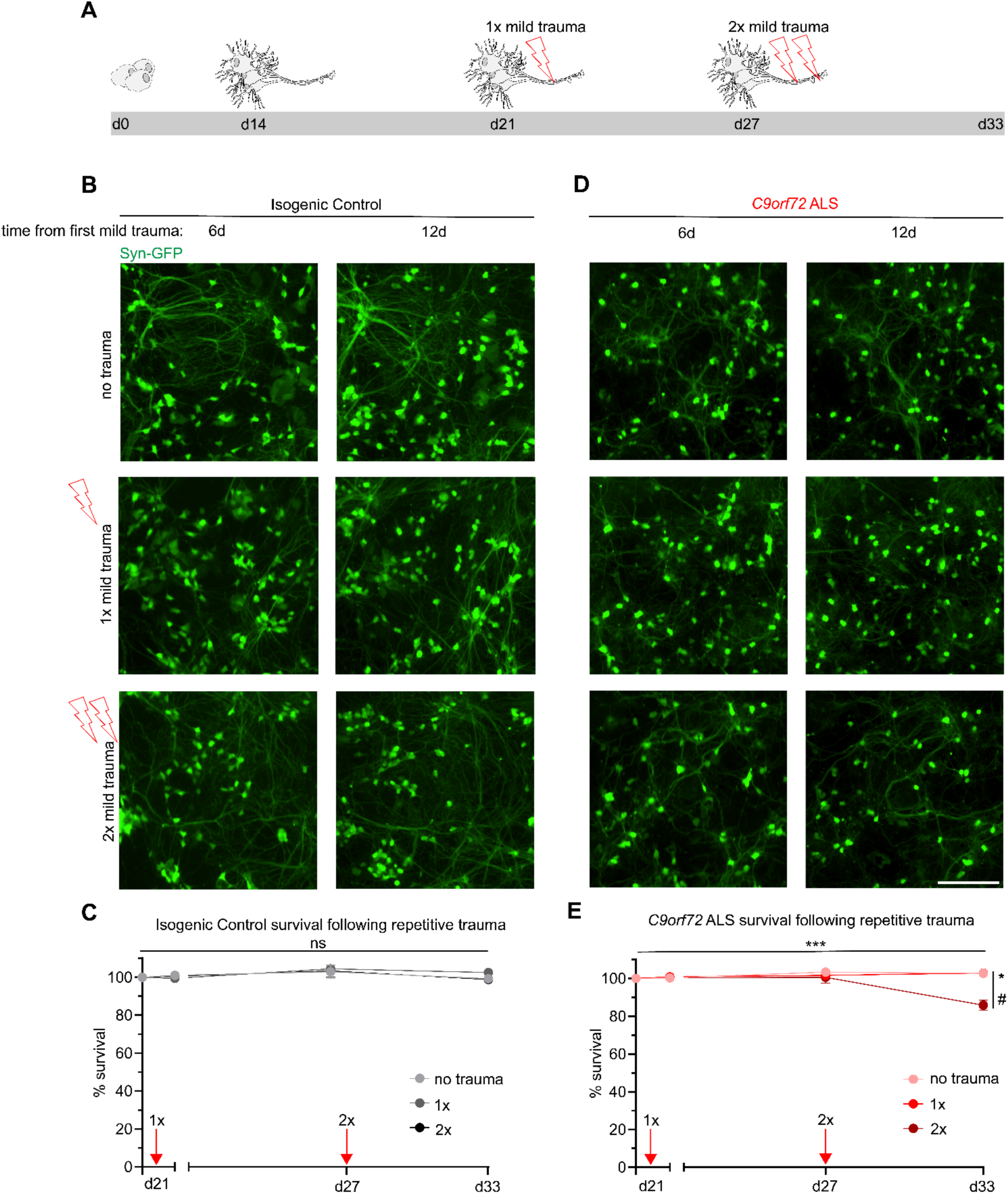
Repetitive mild trauma facilitates neurodegeneration in C9orf72 ALS motor neurons. **(A)** Schematic depicting trauma paradigm for repetitive injury. **(B and D)** Live imaging of CS29 synapsin-GFP motor neurons following either no trauma, 1x mild trauma or 2x mild trauma across an extended timeline (12 days from 1^st^ trauma) in isogenic control (**B**) or *C9orf72* ALS (**D**). Scale bar = 200 μm. **(C and E)** Quantification of synapsin-GFP motor neurons in isogenic control (**C**) or *C9orf72* ALS (**E**) following trauma. Induction of mild trauma is indicated as a red arrow on the graph. Data represent 9 fields of view from 3 independent differentiations, n = 723 – 965 neurons. (**E)** Two-way ANOVA; p < 0.0005. Tukey’s multiple comparisons test 2x vs NTC; * p <0.05 and 2x vs 1x # p < 0.05. Data are represented as mean ± SEM.

**Figure 7.**
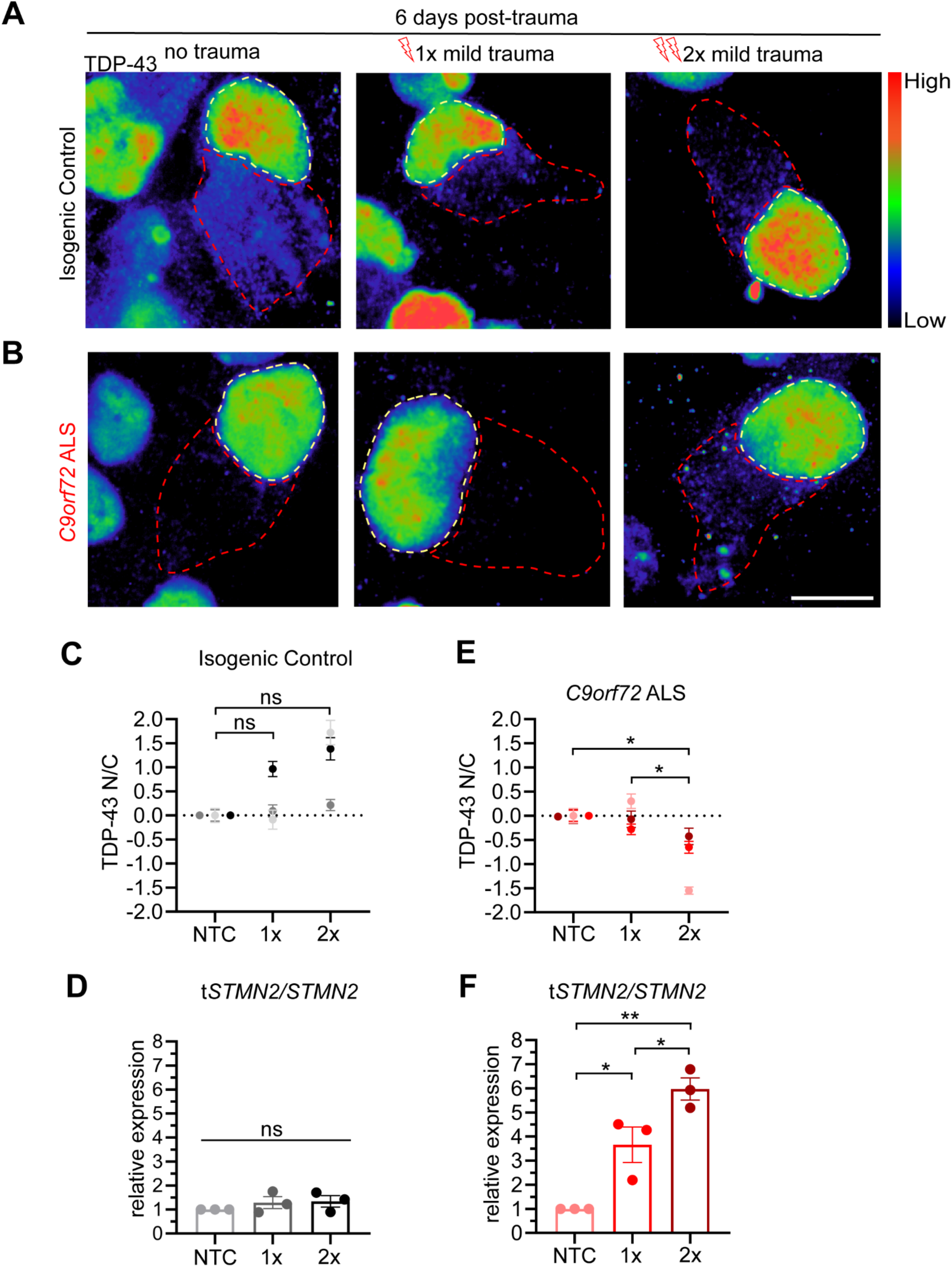
Repetitive mild trauma prevents TDP-43 recovery in C9orf72 ALS motor neurons. (A-B) TDP-43 staining in CS29 neurons 6 days post-trauma following either no trauma, 1x mild trauma, or 2x mild trauma in isogenic control (**A**) or *C9orf72* ALS (**B**) motor neurons. Scale bar = 10 μm. **(C and E)** Quantification of TDP-43 N:C 6 days following mild traumas for either isogenic control (**C**) or *C9orf72* ALS (**E**) motor neurons. (C) One-way ANOVA; NS. (E) One-way ANOVA; p < 0.05. Tukey’s multiple comparisons test relative to NTC indicated with * = p < 0.05; *** = p < 0.0005. 169-184 neurons from 3 independent differentiations. **(D and F)** Quantification of relative expression of truncated STMN2 (tSTMN2) relative to STMN2 6 days post-trauma following either no trauma, 1x mild trauma, or 2x mild trauma for isogenic control (**D**) or *C9orf72* ALS (**F**) motor neurons. 3 biological replicates per condition. (D) One-way ANOVA; NS. (F) One-way ANOVA; p < 0.005. Tukey’s multiple comparisons indicated * = p < 0.05 and ** = p < 0.01. Data are represented as mean ± SEM.

## DISCUSSION

A series of comprehensive epidemiological studies have documented the increased prevalence of ALS in populations exposed to various environmental stressors including traumatic injury [79–82]. However, epidemiological studies are correlational by nature, preventing an explicit link from being drawn. Within this premise, we sought to develop a human iPSC-based preclinical model that would allow us to functionally interrogate the interaction between neurotrauma, degeneration, and associated molecular pathology in the context of patient human ALS MNs. We selected to use MNs with a *C9orf72* HRE on account of the wide range of clinical diversity associated with these patients [18–23], and implemented a custom built mechanical instrument that can deliver biofidelic stretch-based traumatic injury *in vitro* [50–52]. We found that our model faithfully recapitulated core tenets of *C9orf72* ALS pathology in both a transient and persistent manner. *C9orf72* ALS patient MNs were selectively vulnerable to trauma unveiling a distinct susceptibility profile that underscores the complex interplay between genetic predisposition and environmental influences in shaping disease manifestation.

The mechanistic association between ALS and TBI has been investigated using animals including *Drosophila* [69, 83] and rodent models of disease [84–88]. In some of these instances, TBI has been shown to cause stress granule formation, altered N/C transport dynamics, and TDP-43 mis-localization independent of genetic background [49, 68, 69, 84, 88], pathologies that have been documented in ALS patients [41, 43, 44, 89–94]. Moreover, while there are other reports of utilizing iPSC technologies to model TBI mechanisms *in vitro* [95–99] our approach represents the first one to combine ALS patient specific iPSC-derived neurons with a stretch-induced traumatic injury infliction. The HRE in *C9orf72*, which represents the largest genetic cause of ALS and FTD leads to dysfunction and eventual neurodegeneration through both loss-of-function effects as well as gain-of-function RNA and DPR associated toxicity [11–17]. Our findings suggest that the increased vulnerability of *C9orf72* neurons to TBI is primarily mediated by HRE RNA associated toxicity. This is supported by the fact that mild trauma elicited an increase in RNA foci but not poly-GP DPR production even 4 days after injury. This is surprising given that non-canonical translation has been shown to be triggered by the induction of the integrated stress response (ISR) and neuronal hyper-excitability [67], cellular pathways that likely occur downstream of TBI. It is possible that an increase in DPR production would be observed beyond the temporal scope of our analysis or that there is selective translation of non-GP DPRs.

We showed that targeting the C9-HRE with an ASO designed against the sense transcript was partially neuroprotective, implicating the gain-of-function effects of the mutation in the increased susceptibility to severe trauma. The incomplete rescue of the ASO could be related to the contribution of anti-sense *C9orf72* transcript driven toxicity [100, 101] or simply insufficient dosing. Nevertheless, these experiments represent one of the first examples of a pharmacological pre-treatment that can successfully prevent post-traumatic neurodegeneration in human ALS patient neurons.

Our observation that mild trauma led to extended mis-localization of TDP-43 in *C9orf72* ALS MNs, while only a brief mis-localization in control MNs has interesting implications. It suggests that the re-distribution of TDP-43 may be a physiological response to trauma and stress and is in accordance with a series of prior studies. Intriguingly, a recent study used iPSC-derived spheroids from control and mutant *C9orf72* patients and inflicted high-intensity focused ultrasound and similarly found exacerbated TDP-43 dysfunction [99]. TDP-43 has been shown to be associated with cytosolic stress granules following induced stress through chemical treatment [68, 102–104]. Furthermore TDP-43 mis-localization has been observed in animal models of axonal injury, both in cases of TDP-43 ALS-associated mutations and a wild-type genetic background [69, 84, 105, 106]. We propose that the *C9orf72* mutation compromises the typical signaling cascade downstream of injury including the normal re-localization of TDP-43. This could be caused by broad nuclear transport defects that have been previously characterized in both models of ALS [69, 89, 90, 107] and in *Drosophila* models of TBI [69]. Furthermore, the increased mis-splicing of TDP-43 target genes such as *STMN2* even at a time when TDP-43 distribution had recovered to baseline levels, suggests a sustained pathology that extends beyond the initial phase of traumatic injury and could be mediating subsequent vulnerability. This is particularly relevant in the context of repeated mild trauma, which was sufficient to induce neurodegeneration specifically in *C9orf72* ALS MNs. It is possible that nonlethal pathologies such as mis-spliced TDP-43 target genes and C9-HRE nuclear foci accumulate in cells and can ultimately contribute to neurodegeneration.

While the model system we developed here will be a valuable tool as it allows for the investigation of molecular pathology related to traumatic injury in the context of human neurons and a patient-specific genetic background, it also has some major limitations. Firstly, we exclusively focused on the effects of traumatic injury on neurons, however traumatic injury is known to cause non neuronal responses involving oligodendrocytes, and activated astrocytes and microglia [108]. Incorporation of these cell types either directly generated from iPSCs through directed differentiation strategies [109], or in the context of 3D cortical or spinal cord multicellular organoids [110, 111] would allow for a more complete investigation of traumatic injury responses. Secondly, the mechanical injury associated with TBI can come in many ways such as a physical hit, or blast, or even a penetrating injury and downstream of these forces are several tissue damaging mechanisms beyond stretch-induced injury which we modelled here including edema, inflammation etc. [112–115]. Lastly, iPSC-models *in vitro* lack the structural organization and functional connectivity of the intact nervous system in model organisms. At the same time, these limitations can also offer some value in representing a reductionist approach to isolate cell autonomous effects in response to specific types of insults such as stretch-induced injury in this case [116]. Taken together, our results corroborate the value of a preclinical model for traumatic injury, aptly employing stretch injury to simulate physiologically relevant environmental factors. Our work demonstrates the ability to recapitulate clinically relevant pathologies, characterizing the crosstalk between environmental factors and the pathophysiological trajectory of *C9orf72* ALS/FTD.

## ACKNOWLEDGMENTS

We are grateful to the following funding sources: US National Institutes of Health (NIH), National Institute on Neurological Disorders and Stroke (NINDS) and National Institute on Aging (NIA) R01NS104219 (E.K), NIH/NINDS R01NS134166 (E.K), NIH/NINDS R37NS057553 and RF1NS101986 (F-B.G), NIH/NINDS R01NS113935 (J.D.F), the US Department of Defense/National Defense Education Program through the Educational and Research Training Collaborative at the University of Illinois Chicago, Award HQ00342010037 (A.M and C.S), the Patrick Grange Memorial Foundation (E.K and J.D.F), the Les Turner ALS Foundation (E.K) and the New York Stem Cell Foundation (E.K). E.K is a Les Turner ALS Center Investigator and a New York Stem Cell Foundation – Robertson Investigator.

## AUTHOR CONTRIBUTIONS

Conceptualization: E.J.M, E.K; Methodology: E.J.M, C.S, G.E, A.M, I.A.E, G.K, E.K; Investigation: E.J.M, C.S, A.M, I.A.E, G.K; Writing – Original Draft: E.J.M, E.K; Writing – Review & Editing: E.J.M, C.S, G.E, A.M, I.A.E, G.K, FG.B, J.D.F and E.K; Funding Acquisition: FG.B, J.D.F and E.K; Supervision: FG.B, J.D.F and E.K.

## MATERIALS AND METHODS

### Stem cell culture

Human induced pluripotent stem cells (hiPSC) *C9orf72* mutant and isogenic control pair (CS29 and CS52) were obtained from the Cedars Sinai iPSC repository. Isogenic controls were generated by CRISPR/Cas9 gene editing targeting the regions immediately 5’ and 3’ of the repeat expansion. Stem cells were maintained on Matrigel (BD Biosciences) with mTeSR1 media (Stem Cell Technologies) and passaged using Accutase (Innovative Cell Technologies). All cell cultures were maintained at 37°C and 5% CO2. All iPSC cultures are tested regularly for mycoplasma. All lines used were mycoplasma-free. Passaged iPSC cultures were kept in Y27632 for 24 hours (DNSK international).

### Motor neuron differentiation

When hiPSC colonies were ∼80% confluent, media was replaced with N2B27 medium (50% DMEM:F12, 50% Neurobasal, supplemented with NEAA, Glutamax, N2 and B27; Thermo Fisher Scientific) containing 10μM SB431542 (DNSK International), 100nM LDN-193189 (Bio-Techne), 1 μM Retinoic Acid (RA, Millipore-Sigma) and 1μM of Smoothened-Agonist (SAG, DNSK International). Media was changed daily. On day 6, culture media was switched to N2B27 medium supplemented with 1μM RA, 1μM SAG, 5μM DAPT (Bio-Techne) and 4μM SU5402 (DNSK International), fed daily until 14 total days of differentiation. On day 14, MNs were dissociated using TrypLE Express (Thermo Fisher Scientific) supplemented with DNase I (Worthington) and plated onto murine astrocytes seeded on a polydimethylsiloxane (PDMS) plate (Nunc) pre-coated with Poly-D-Lysine (BD Biosciences). MNs were cultured in Neurobasal medium supplemented with NEAA, Glutamax, N2, B27, Ascorbic acid (0.2 μg/ml; Sigma-Aldrich) BDNF, CNTF and GDNF (10ng/mL, Bio-Techne), 2% FBS (Gemini), and P/S (Thermo Fisher Scientific). For the first 24 hours, MNs were maintained in Y. Media was half-changed every 2-3 days after the first 24 hours.

### Simulation of stretch trauma

Prior to experimentation, a high-speed camera was used to establish a zero plane for PDMS plates, defined as the point where the posts for injury encountered the PDMS bottom but did not cause displacement in the membrane. Displacements and injuries were performed using custom software (Python) controlling the post array via a servo controller. Plates were clamped down using thick acrylic to hold the plates stationary during injury in a manner that would not contribute additional stress to the plates (the force from securing the plates was eventy distributed across the plate). Prior to injury, posts were lubricated using vegetable oil (Kroger) to minimize friction and adhesion between the posts and the PDMS during trauma. Trauma simulations spanned a period of ∼70 msec. Displacements were selected based on previously calculated quantifications of Lagrangian strain.

### Live imaging of *SYN1*-GFP cells

For live imaging experiments, lentiviral *SYN1*-GFP (PZ196, Addgene) was added to neurons in suspension following dissociation on day 14. Neurons were plated in culture media with *SYN1*-GFP for the first 24 hours and following a full media change were cultured as previously described. For nuclear staining, NucSpot Live 650 Nuclear Stain (Biotium) was added 1:1000 to media 1 hour prior to imaging and supplemented with media changes. Propidium iodide (Sigma) was added to media 1:2000 and supplemented with media changes. Images were collected prior to trauma and every 1.5 hours for the first 6 hours following trauma. After 6 hours, images were collected every 12 hours. Images showing synapsin-GFP alone were collected from an IncuCyte S3 (Sartorius). Images showing PI and nuclear stains were collected using a Nikon BioStation (Nikon). Image Analysis was performed using Image J (U. S. National Institutes of Health, Bethesda, Maryland, USA)

### Antisense oligonucleotide treatment

Following MN dissociation, ASOs were added to day 14 motor neurons in suspension at a concentration of 5μM (C9orf72: CCGGCCCCGGCCCCGGCCCC; Control: CCUUCCCTGAAGGTTCCUCC; Integrated DNA Technologies). Every 3 days, media was supplemented with 2.5μM ASOs. On the 10^th^ day, RNA was collected for qRT-PCR. Trauma experiments were performed following 10 days of ASO administration.

### RNA preparation and qRT-PCR

Cells were harvested by direct lysis in TRIzol Reagent (Life Technologies). RNA isolation was performed via phase separation according to manufacturers’ protocol. 1-2μg of RNA was used as input for generation of cDNA according to the SuperScript IV First-Strand cDNA Synthesis Reaction Kit (Thermo Fisher Scientific). RT-PCR was performed using PowerUp SYBR Green Master Mix (Thermo Fisher Scientific) on the CFX system (Bio-Rad). Two housekeeping genes (RPLP0/CYC1) were averaged for determination of the cycle of threshold (Ct). For ASO effectiveness, this Ct was subtracted from the average Ct for *C9orf72* to obtain the ΔCt. Relative gene expression was calculated as 2−ΔCt (ΔΔCt) and expressed relative to the scrambled ASO treatment. For splicing experiments, the average Ct for the gene of interest was subtracted from the average Ct for the spliced isoform for a determination of splice:total expression.

### List of qPCR primers

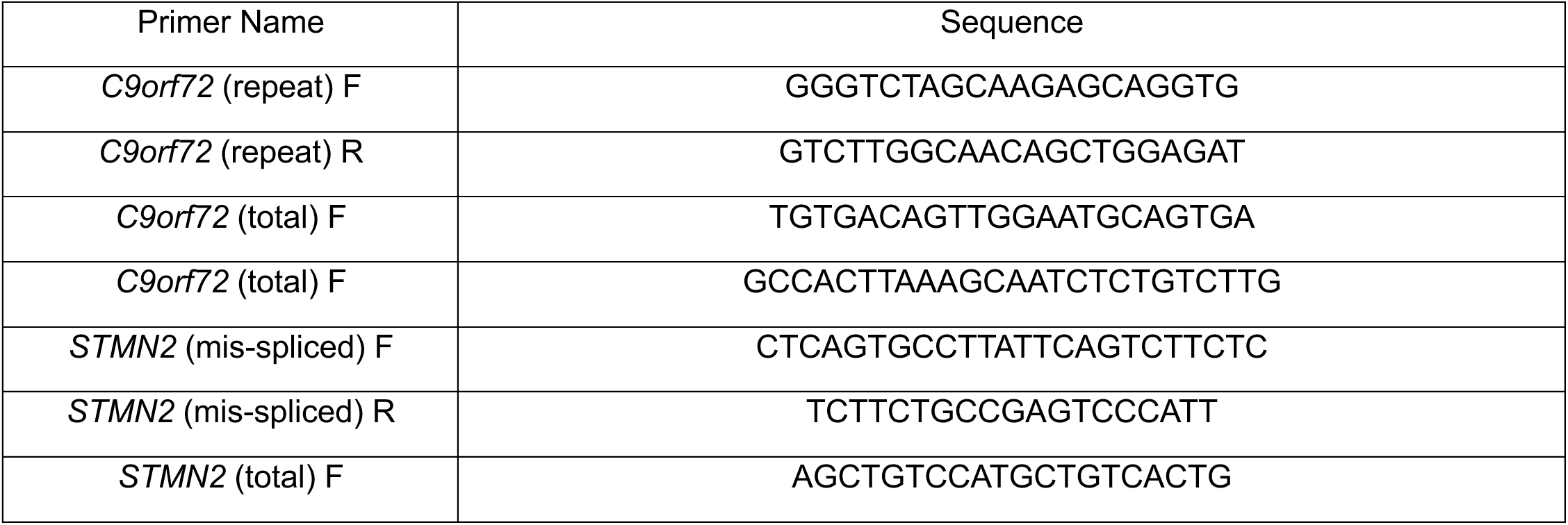
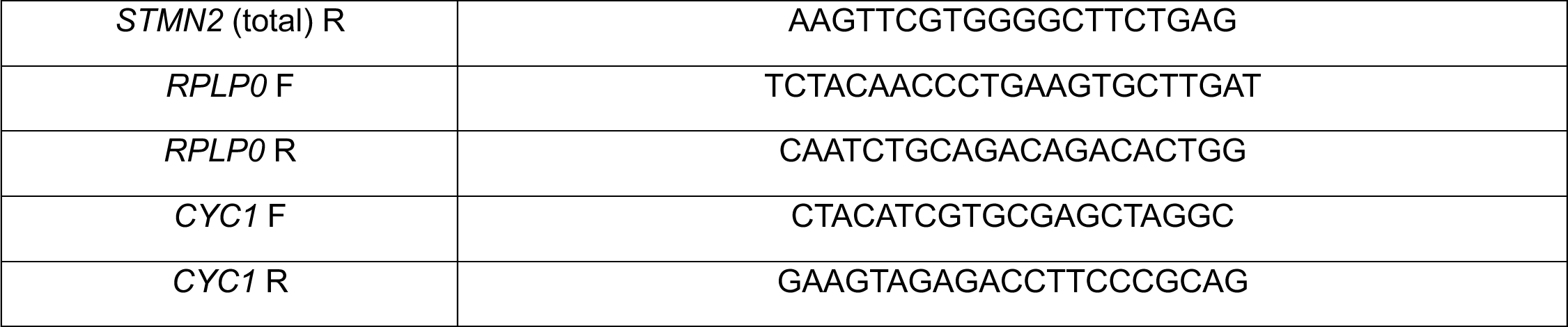

### Fluorescent in situ hybridization (FISH)

FISH was performed as previously detailed [117]. Briefly, cells were cultured in PDMS plates, fixed with 4% PFA, and permeabilized with 0.3% Triton for 15 minutes. Prior to hybridization, cells were equilibrated in 50% formamide (Spectrum) + 2X SSC (Invitrogen) for 10 minutes at 60°C. During equilibration, probe mixture was preheated at 95°C for 10 minutes. To hybridize, 27 μL of probe mixture in 200 μL of hybridization buffer were added to each well, and cells were incubated at 60°C for 1 hour. Following hybridization, cells were incubated with 50% formamide + 2X SSC at 65°C for 25 minutes, then with fresh 50% formamide + 2X SSC at 65°C for 15 minutes, and finally with 1X SSC + 40% formamide at 60°C for 10 minutes. Cells then underwent a series of room temperature washes: 3x with 1X SSC, 2x 5 minutes with 1X SSC, 1x 5 minutes with Tris-buffered saline (Invitrogen), and finally 1x 5 minutes with Tris-glycine buffer (Invitrogen). Cells were immersed in 3% PFA for post-fixation and then incubated for 1 hour with blocking buffer at room temperature. Cells were incubated in immunofluorescence buffer with primary antibodies at 4°C overnight. Following a brief wash in immunofluorescence buffer, cells were placed in immunofluorescence buffer with secondary antibodies for 30 minutes at room temperature. Following incubation with secondary antibodies, cells were sequentially washed with immunofluorescence buffer, Tris-buffered saline, Tris-glycine buffer, 2mM MgCl2 in PBS, and PBS. DAPI (Invitrogen) was added to wells in PBS and imaging was performed immediately after.

Hybridization buffer consisted of 40% formamide, 1mM ribonucleoside vanadyl complex (Sigma-Aldrich), 10mM NaPO4, BSA (2mg/ml, Roche), and 1X SSC. Probe mixture for detection of C9 RNA foci consisted of 25 μL 80% formamide, 0.5 μL E. coli tRNA (20 μg μL, Thermo Fisher Scientific), 1 μL salmon sperm (10 μg/ μL, Thermo Fisher Scientific), and 0.4 μL 25μM 5′ digoxigenin (DIG)-labeled CCCCGGCCCCGGCCCC locked nucleic acid probe (Exiqon, batch 620574). Tris-buffered saline was comprised of 200mM Tris solution, pH 7.4, 50mM Tris and 15mM NaCl solution, pH 7.4 and 0.75% glycine. Blocking buffer consisted of 5% heat-shocked BSA in Tris-buffered saline and 10% normal donkey serum (Jackson Immunoresearch). Immunofluorescence buffer was comprised of 2% heat-shocked BSA in Tris-buffered saline. Detection of DIG-labeled probe was performed with a fluorescein-conjugated sheep anti-DIG antibody (1:250, Roche). All buffers were made with RNase-free PBS or water.

### Quantification of C9-GP DPR by ELISA

Cell pellets of *C9orf72* and isogenic control MNs co-cultured with murine astrocytes were collected and lysed in RIPA buffer (Thermo Fisher Scientific). Lysed material was sonicated for 5 pulses of 40Hz. Lysed material was centrifuged at 22,000 rpm for 15 minutes and supernatant was transferred to a new tube. The concentration of protein in the supernatant was quantified using the BCA protein assay (Thermo Fisher Scientific). 400µg of protein lysate was used for quantification of poly-GP. Lysates were loaded onto a 96-well plate (Meso Scale Discovery, Cat. no. L45XA) that had been coated with a custom-made polyclonal anti-(GP)_8_ antibody (1 µg/ml, Covance). Lysates were compared in multiple wells. Serial dilutions of recombinant (GP)8 peptide were prepared in 1% BSA+TBS-T for generation of the standard curve. For detection, anti-GP(8) tagged with GOLD SULFO (GOLD SULFO-TAG NHS-Ester Conjugation Pack, Meso Scale Discovery; Cat. no. R31AA) was used at a concentration of 0.5µg/mL. Quantification of signal from the assay plate was performed using a QuickPlex SQ120 instrument (Meso Scale Discovery). Sample groups were blinded prior to analysis.

### Immunocytochemistry

Fixation of cells was performed with 4% paraformaldehyde. Subsequently, cells were blocked in PBS with 0.2% Triton (Sigma-Aldrich) and 10% normal donkey serum (Jackson ImmunoResearch). Samples were incubated at 4°C overnight with primary antibodies in PBS with 5% wt/vol BSA. Following 3 washes with PBS + 0.1% Triton, samples were incubated at room temperature for 1 h with corresponding secondary antibodies conjugated to Alexa Fluor 488, Alexa Fluor 555, and Alexa Fluor 647. Finally, cell nuclei were stained with Hoechst 33342. Immunolabeled samples were imaged using a Nikon Ti2 Widefield and W1 spinning disk confocal photographed using a C10600-ORCA-R2 digital CCD camera (Hamamatsu Photonics, Japan).

### List of immunofluorescence reagents

List of immunofluorescence reagents:
| Reagent (host) | Dilution | Source (catalogue number) |
| --- | --- | --- |
| Hoechst | 1:200 | Invitrogen H3570 |
| ISL1/2 (mouse) | 1:100 | Development Studies Hybridoma Bank, University of Iowa, AB_2314683 |
| TDP-43 (rabbit) | 1:200 | ProteinTech 10782-2-AP |
| MAP2 (chicken) | 1:1000 | Abcam ab5392 |
| Alexa 488 anti-mouse (donkey) | 1:500 | Invitrogen A21202 |
| Alexa 488 anti-rabbit (donkey) | 1:500 | Invitrogen A21206 |
| Alexa 555 anti-mouse (donkey) | 1:500 | Invitrogen A31570 |
| Alexa 555 anti-rabbit (donkey) | 1:500 | Invitrogen A31572 |
| Alexa 647 anti-chicken (donkey) | 1:500 | Jackson Immuno Research Labs 703-606-155 |

### Image acquisition and analysis

Quantification of *SYN1*-GFP and propidium iodide was performed using ImageJ (U. S. National Institutes of Health, Bethesda, Maryland, USA) with the cell counter analysis plugin. In Figure 1, images were collected within 1.5 hours prior to trauma and then every 1.5 hours for the first 6 hours following trauma. Beyond this, images were collected every 12 hours for a total of 54 hours. For Figure 2, images were collected and *SYN1*-GFP quantified 1.5 hours prior to and 1.5 hours post-trauma. Images used for analysis in Figures 1, 2, and 6 were obtained using the IncuCyte S3 live cell imager at 20X magnification with an exposure time of 300ms. For Figure 3, quantitative analysis of RNA foci using FISH was performed on a Nikon W1 Spinning Disk Confocal using a 60x objective lens (slice depth 0.6µm). For Figures 4, 5, 7, and the accompanying Supplemental Figures analysis of TDP-43 staining was performed using images gathered from a Nikon Ti2 Widefield microscope at 40X magnification (slice depth = 1.2 µm; exposure times for DNA: 200ms, TDP-43: 800ms, MAP2: 2s). Representative images for TDP-43 localization analysis were obtained using a Nikon W1 Spinning Disk Confocal with a 100x silicone oil objective lens (slice depth = 0.4µm). For Figure 6, images of *SYN1*-GFP neurons were taken immediately prior to injury and at 6 and 12 days later at 20X with an exposure time of 300ms using the IncuCyte S3 live cell imager.

All experiments consisted of 3 independent differentiations, with at least 2 biological replicates per condition per experiment. Images for data analysis consisted of at least 9 randomly selected optical fields. For Figures 1, 2, and 6 *SYN1*-GFP was quantified by creating a threshold for *SYN1*-GFP positive neurons using Fiji. Figure 3 was analyzed using Nikon Elements software. Quantification of RNA foci was performed using a gain of 1.5 and background thresholded out on a per-slice basis. For Figures 4, 5, and 7 image analysis was performed using Nikon Elements analysis software. Analysis of nucleocytoplasmic ratios was performed using ROI tracing to compartmentalize the cytosol and nucleus. ROI analysis was used to determine the average (mean) fluorescence intensity across the region for both nuclear and cytosolic compartments, and the ratio represented was calculated by dividing the average nuclear fluorescence intensity by the average cytosolic fluorescence intensity. Experimental averages across differentiations for Figures 4C, 4E, 5B, 5D, 5F, 5H, 7C and 7E were standardized to 0 through z-score conversion. Data points were calculated relative to the experimental average and standard deviation based on the “no trauma” control condition of each respective experiment. Individual values were then standardized based on the degree of deviation from the experimental average relative to the experimental standard deviation. Images were analyzed and assembled as maximum projections from z-stack slices at the depths stated above. Representative images are presented with the same thresholding within each experiment. Images were assembled in Adobe Illustrator following direct export from Nikon Elements software. All experimental conditions were blinded to the operator prior to analysis.

### Statistical analysis

All statistical analysis was performed using GraphPad Prism 10 software. Data from multiple experiments were normalized using z score conversion, which standardizes the experimental means to 0 and experimental standard deviation to 1 for each independent experiment. Baseline data were then validated to be normal using the Anderson-Darling test.

Figure 1G: Survival analysis was performed using a two-way ANOVA with Tukey’s post-hoc multiple comparisons test indicating statistical significance between experimental group.

Figure 2C: qPCR analysis of ASO knockdown was performed in 3 biological replicates, each in triplicate. A mean of these 3 replicates is represented.

Figure 2D: analysis of *SYN1*-GFP quantification was performed using two-way ANOVA with Tukey’s post-hoc multiple comparisons tests indicating statistical significance between groups.

Figure 3: two-way ANOVA was performed for panels C, F, and G. Experimental averages from 3 independent experiments are represented for RNA foci. For DPR analysis, data are represented as the mean of 3 independent experiments performed concurrently. Tukey’s post-hoc multiple comparisons test indicates statistical significance between groups.

Figure 4C and 4D: statistical analysis was performed using a one-way ANOVA, with a Tukey’s post-hoc multiple comparisons test indicating statistical significance between groups.

Figure 5B, 5D, 5F, and 5H: statistical analysis was performed using a one-way ANOVA, with a Tukey’s post-hoc multiple comparisons test indicating statistical significance between groups.

Figure 5J and 5K: statistical analysis was performed using a Welch’s t-test.

Figure 6C and 6E: statistical analysis was performed using a two-way ANOVA, with a Tukey’s post-hoc multiple comparisons test indicates statistical significance between groups.

Figure 7C-F: statistical analysis was performed using a one-way ANOVA, with a Tukey’s post-hoc multiple comparisons test indicates statistical significance between groups.

### Justification of statistical tests

Z scores were measured to conservatively assess interexperimental conditions based on a standardized normalization procedure that allows both standard deviation and means to be the same across experiments (mean = 0, 1 standard deviation = 1). Interexperimental variations were determined to be similar enough to use this method of normalization. Two-way ANOVA tests were used for survival analysis due to the movement of neurons into and out of frame, which could present confounding data if presented as a Kaplan-Meyer curve. In many cases, there could not be explicit tracing of a specific neuron. A multitude of frames and hundreds of cells were analyzed for an accurate representation of culture, and two-way ANOVA was used because of variables for treatment conditions x time. For stretch experiments, a one-way ANOVA with Tukey’s multiple comparison post-hoc correction was used to determine statistical significance. Statistics were performed on one line where the only variable was treatment condition. Experimental averages are shown with standard error, and each experimental condition ranged 40-75 cells to gather an accurate assessment of the population.

## FIGURES AND FIGURE LEGENDS

**Supplementary Figure 1.**
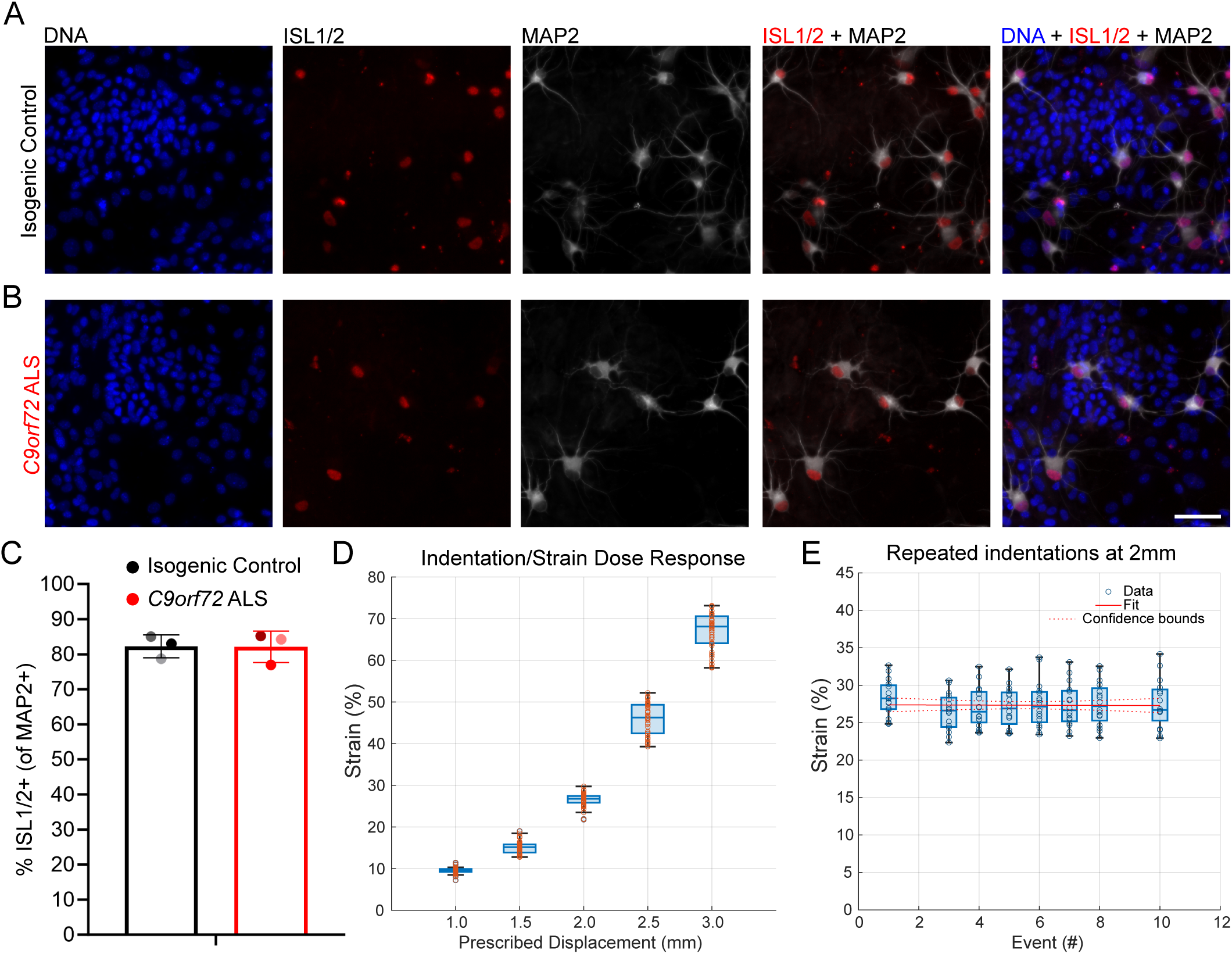
Efficiency of differentiation in CS29 isogenic control and C9orf72 ALS. **(A)** Immunofluorescence depicting DNA, ISL1/2, and MAP2 in CS29 isogenic control and (**B)** *C9orf72* ALS. Scale bar = 50 μm. **(C)** Graphical representation of experimental averages quantifying % ISL1/2+ motor neurons among MAP2+ neurons from 3 independent differentiations. Analysis performed with 24-28 fields of view quantifying 621-628 MAP2+ neurons. Mann Whitney p = NS. Data are represented as mean ± SEM. **(D)** The effect of increasing the nominal indentation depth on the membrane strain. **(E)** The evolution of the membrane strain over the course of repeated indentations procedures (red line = linear regression with slope = -0.06±0.08 p = 0.443).

**Supplementary Figure 2.**
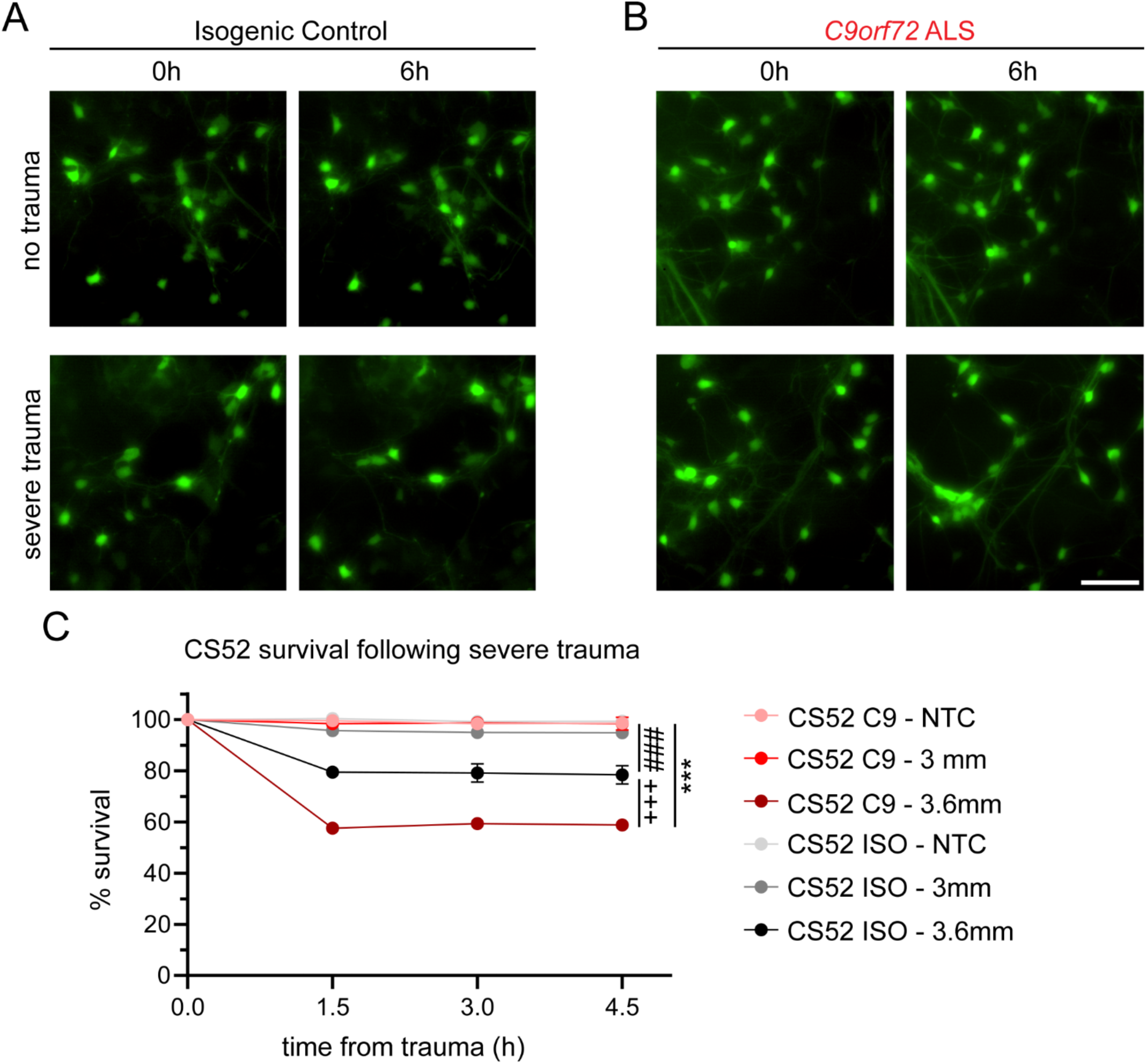
C9orf72 ALS selective vulnerability is observed in the CS52 cell line. **(A)** Live imaging showing CS52 synapsin-GFP transduced motor neurons before and after either no trauma or 3.6mm trauma in isogenic control or **(B)** *C9orf72* ALS. Scale bar = 50 μm **(C)** Quantification of synapsin-GFP motor neurons following trauma. Two-way ANOVA p < 0.0005 from 2-4 biological replicates of 3 independent differentiations comprising 280-656 motor neurons. Tukey’s multiple comparison *** = p < 0.0005 vs C9 NTC, ### = p < 0.0005 vs ISO NTC, +++ = p < 0.0005 vs ISO 3.6mm. Data are represented as mean ± SEM.

**Supplementary Figure 3.**
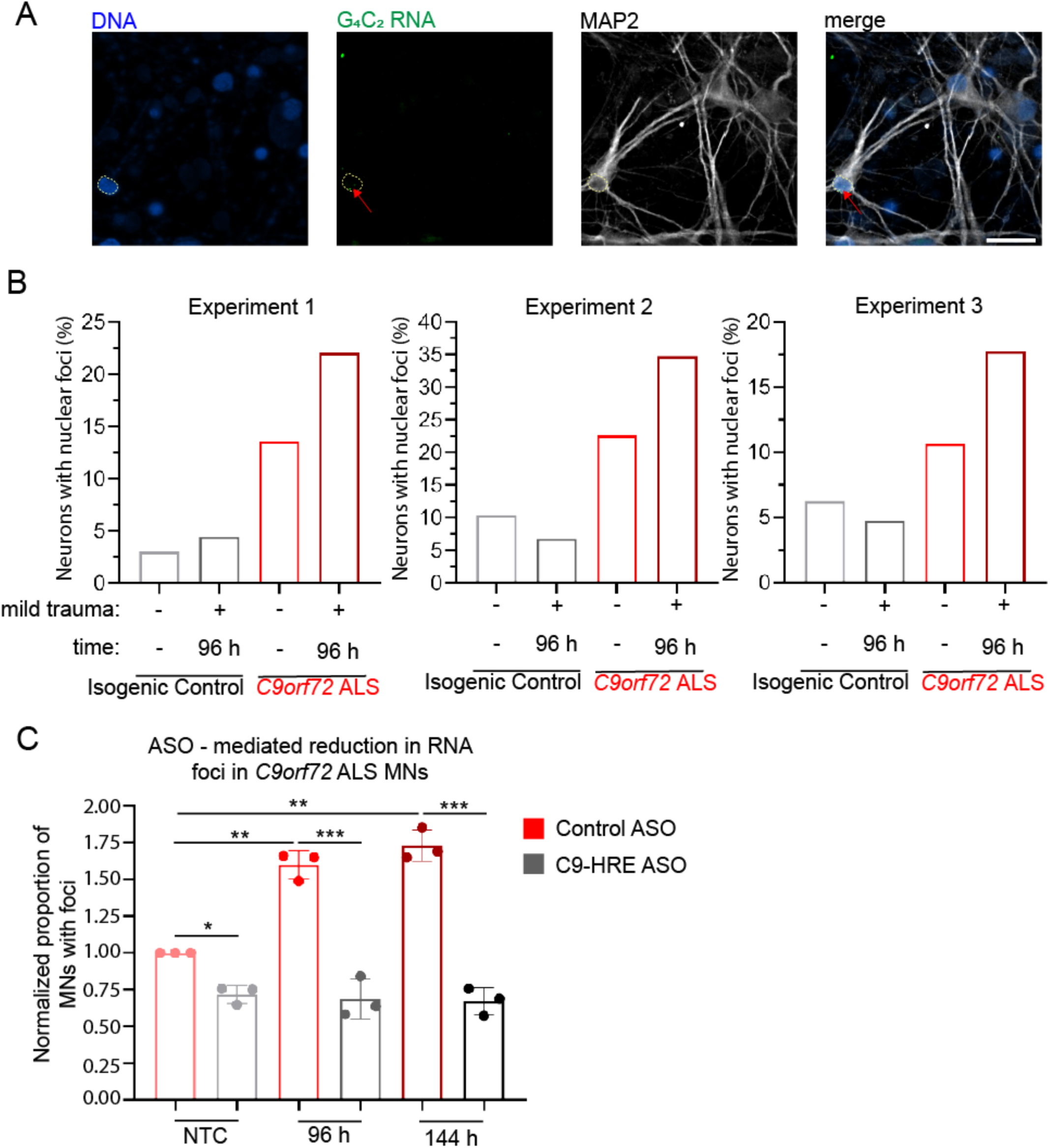
Single channel characterization and effect of ASO pretreatment on nuclear G4C2 foci using FISH. (A) Single channel zoomed out visualization of image used in Figure 3E. Red arrow indicates RNA foci and circle shows nuclear area. Scale bar = 25 μm. (B) Experimental quantification of 3 independent differentiations noting the percentage of neurons with nuclear RNA foci. (C) Experimental data quantifying the effects of ASO pretreatment on RNA foci prevalence. Data is presented as normalized proportion of MNs with C9orf72 RNA foci relative to no trauma control (NTC) condition treated with control ASO. Data are collected from 3 independent differentiations quantifying 145-197 neurons. Two-way ANOVA p < 0.0005. Tukey’s multiple comparison *** = p < 0.005, ** = p < 0.01, * = p < 0.05 relative to *C9orf72* ALS + control ASO. Data are represented as mean ± SEM.

**Supplementary Figure 4.**
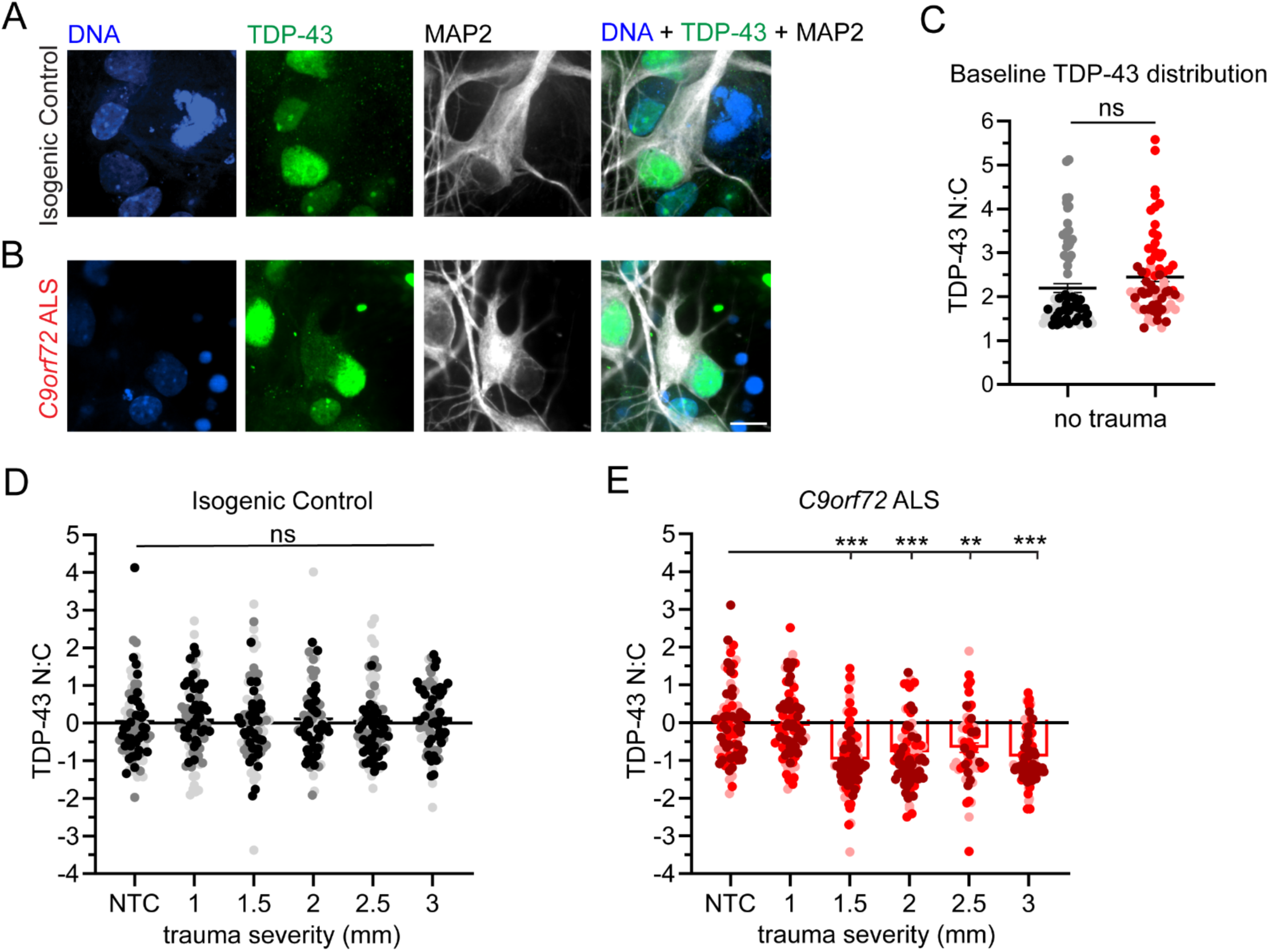
Single channel characterization of TDP-43 staining 4 hours post-trauma. (A-B) Single channel visualization of TDP-43 distribution in motor neurons in either isogenic control (**A**) or *C9orf72* ALS (**B**). Scale bar = 10 μm. **(C)** Graphical representation of baseline TDP-43 distribution used in Figure 4. Different shades represent experimental groups for either isogenic control (82) or *C9orf72* ALS (75) motor neurons. **(D-E)** Individual quantification of TDP-43 N/C across 3 independent differentiations in isogenic control (**D**) or *C9orf72* ALS (**E**) motor neurons. Different shades represent experimental distributions of individual cells. (**D**) Two-way ANOVA NS; (**F)** Two-way ANOVA p < 0.0005. Tukey’s multiple comparisons test vs NTC. ** = p < 0.01, *** = p < 0.005. Data are represented as mean ± SEM.

**Supplementary Figure 5.**
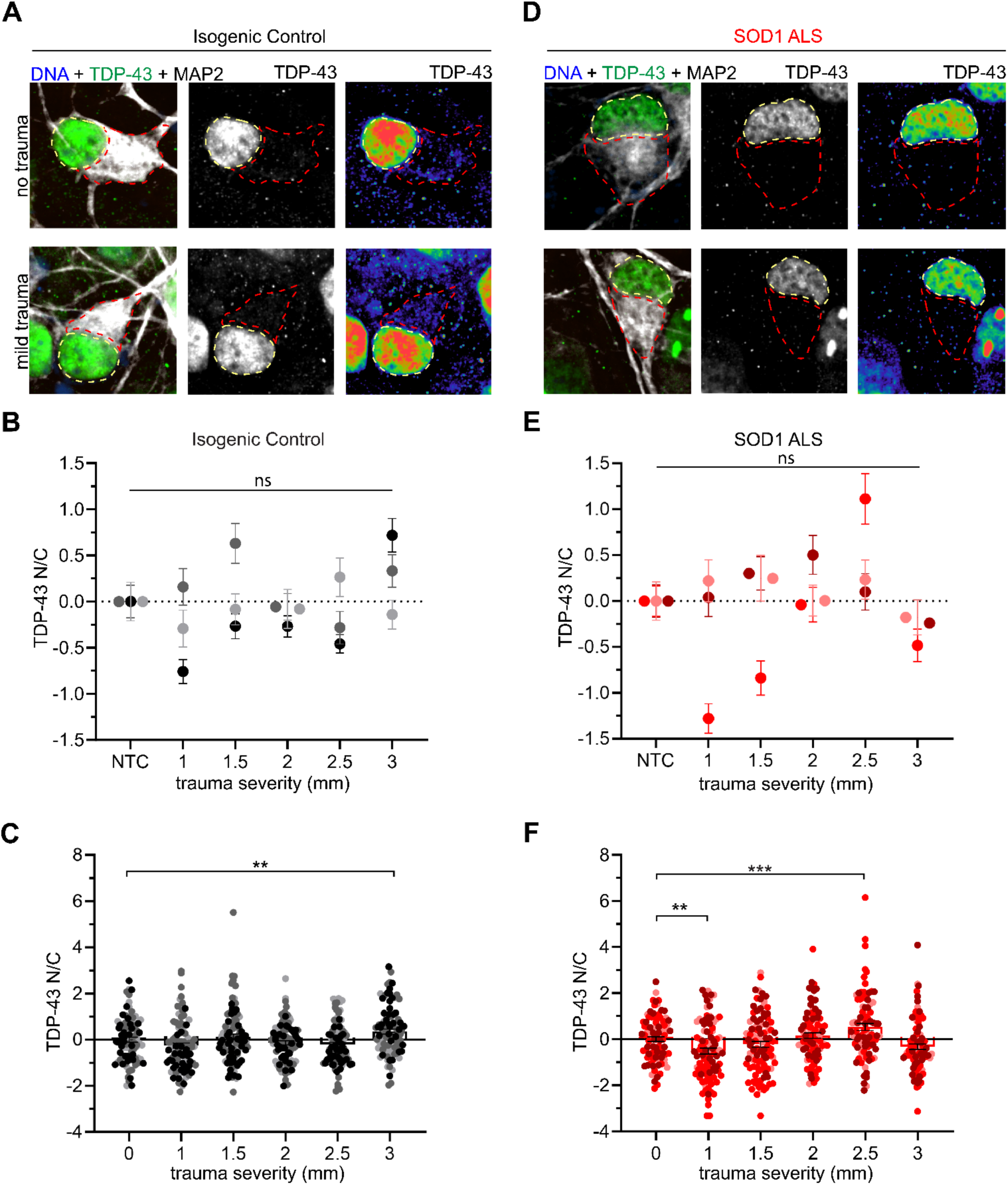
SOD1 ALS motor neurons do not show cytosolic accumulation of TDP-43 following trauma. (A and. **D)** Immunocytochemistry showing TDP-43 distribution in motor neurons under no trauma and mild trauma conditions in isogenic control (**A**) and SOD1 ALS (**D**). Scale bar = 10 μm. **(B and E)** Quantification of experimental averages from 3 independent differentiations of TDP-43 nucleocytoplasmic ratios in isogenic control (**C**) or SOD1 (**D**) motor neurons. One-way ANOVA, NS. **(C and F)** Individual data points corresponding to one TDP-43 nucleocytoplasmic ratio per cell showing the overall data distribution in isogenic control (**C**) or SOD1 ALS (**F**) motor neurons. Analysis was performed on n = 87 - 104 cells per condition. One-way ANOVA, p < 0.0001. Tukey’s multiple comparison test relative to NTC is indicated. ** = p < 0.01 *** = p < 0.005. Data are represented as mean ± SEM.

**Supplementary Figure 6.**
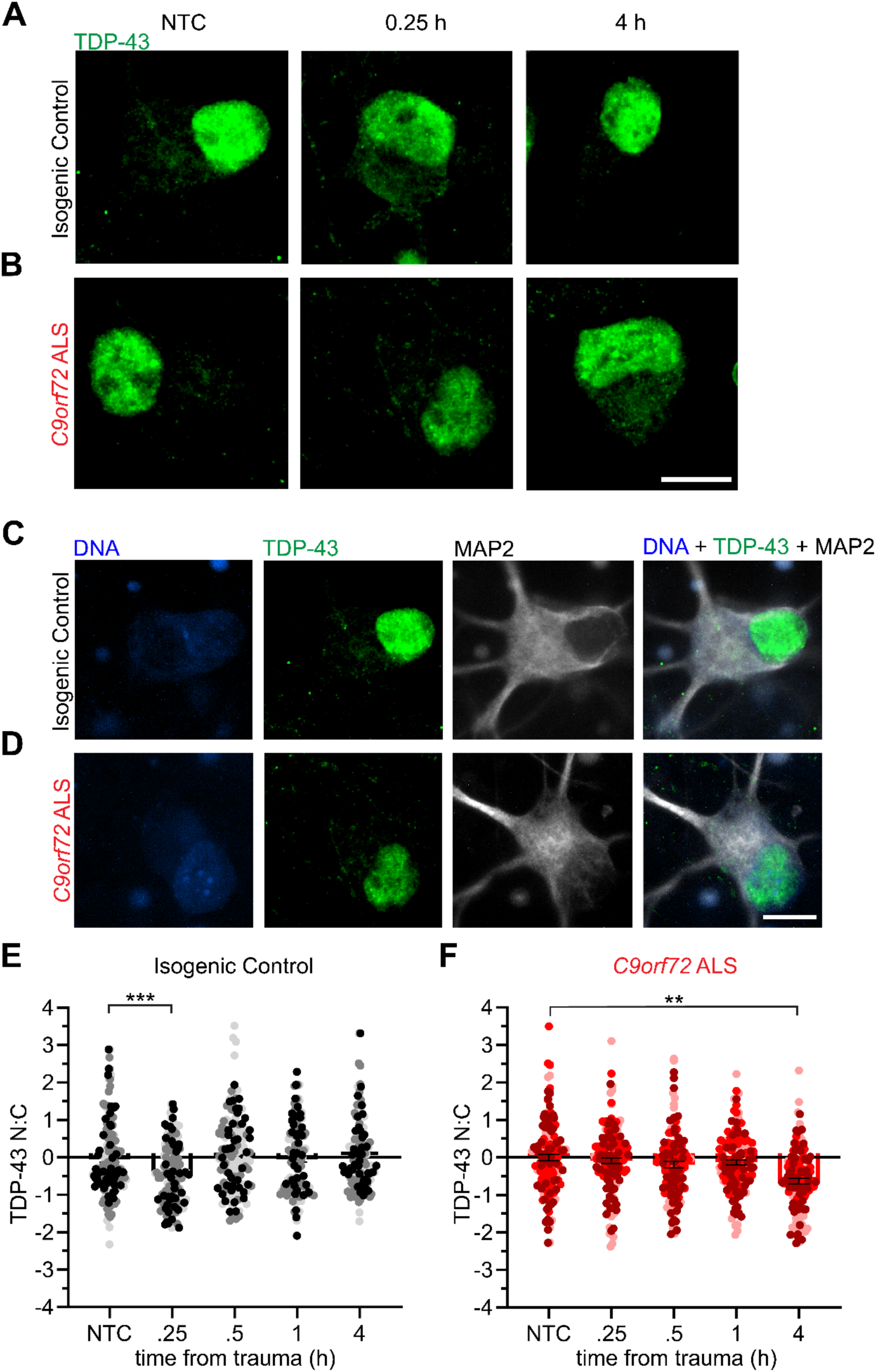
Single channel characterization and individual data point analysis of motor neurons assessing immediate TDP-43 distribution following mild trauma. **(A**-**B)** Visualization of representative GFP (TDP-43) staining without heatmap filter in isogenic control (**A**) or *C9orf72* ALS (**B**) motor neurons. Scale bar = 10 μ. **(C-D)** Single channel visualization of representative images from either isogenic control (**C**) or *C9orf72* ALS (**D**) motor neurons. Scale bar = 10 μm. **(E-F)** Data from individual cells used for quantification of experimental averages in Figure 5 for either isogenic control (**E**) or *C9orf72* ALS (**F**). Data are motor neurons from 3 independent differentiations. Different shades represent different experiments. n = 157-193. **(E)** One-way ANOVA, p < 0.05; **(F)** p < .0001; Tukey’s multiple comparison test relative to NTC is indicated; ** = p < 0.01 *** = p < 0.005.

**Supplementary Figure 7.**
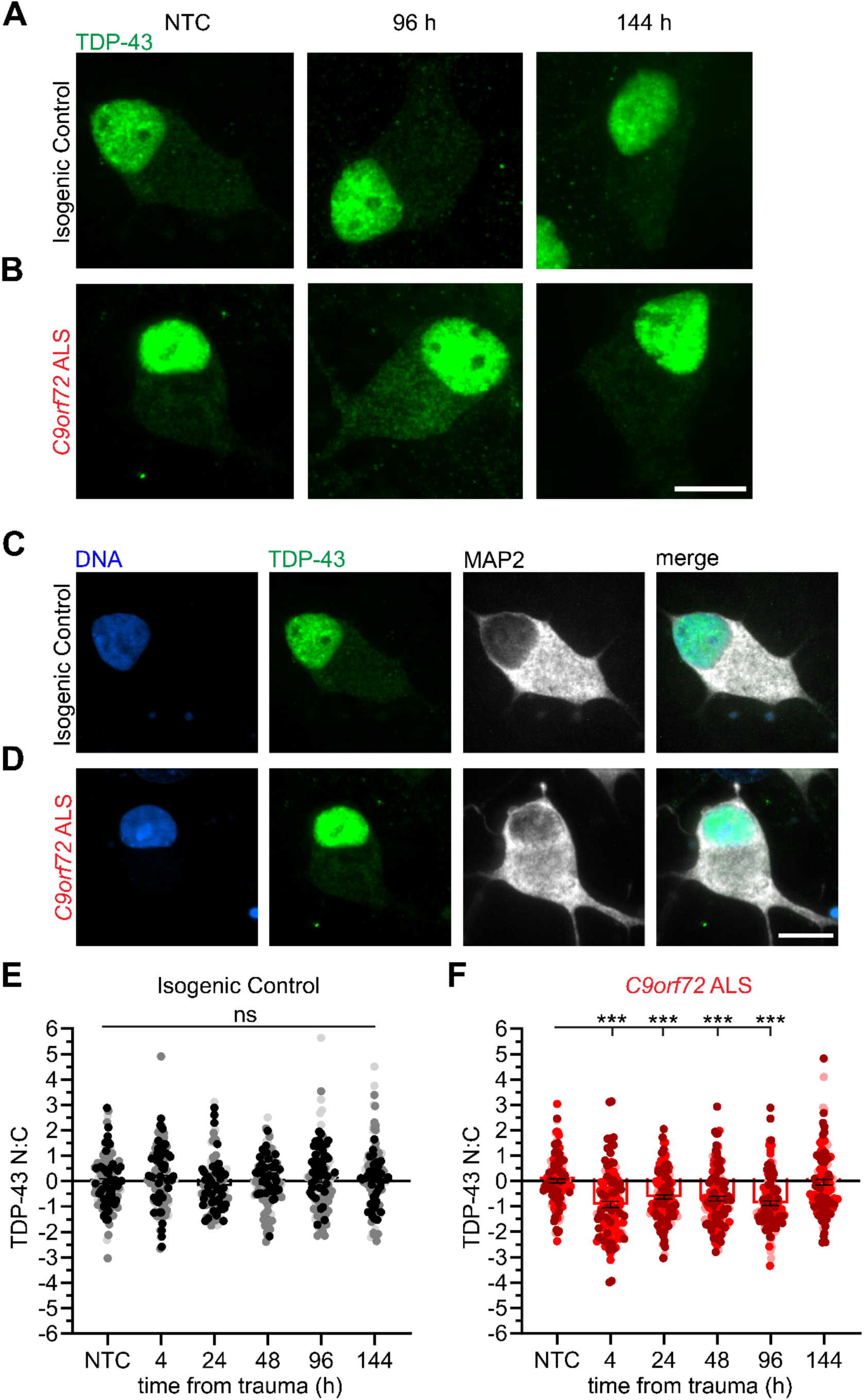
Single channel characterization and individual data point analysis of motor neurons assessing extended TDP-43 distribution following mild trauma. (A-B) Visualization of representative GFP (TDP-43) staining without heatmap filter in isogenic control (**A**) or *C9orf72* ALS (**B**) motor neurons. Scale bar = 10 μm. **(C-D)** Single channel visualization of representative images from either isogenic control (**C**) or *C9orf72* ALS (**D**) motor neurons. Scale bar = 10 μm. **(E-F)** Data from individual cells used for quantification of experimental averages in Figure 5 for either isogenic control (**E**) or *C9orf72* ALS (**F**). Data are motor neurons from 3 independent differentiations. Different shades represent different experiments. n = 124-133. One-way ANOVA, NS for **E**; p < 0.0001 for **F**. Tukey’s multiple comparison test relative to NTC is indicated; *** = p < 0.0005.

**Supplementary Figure 8.**
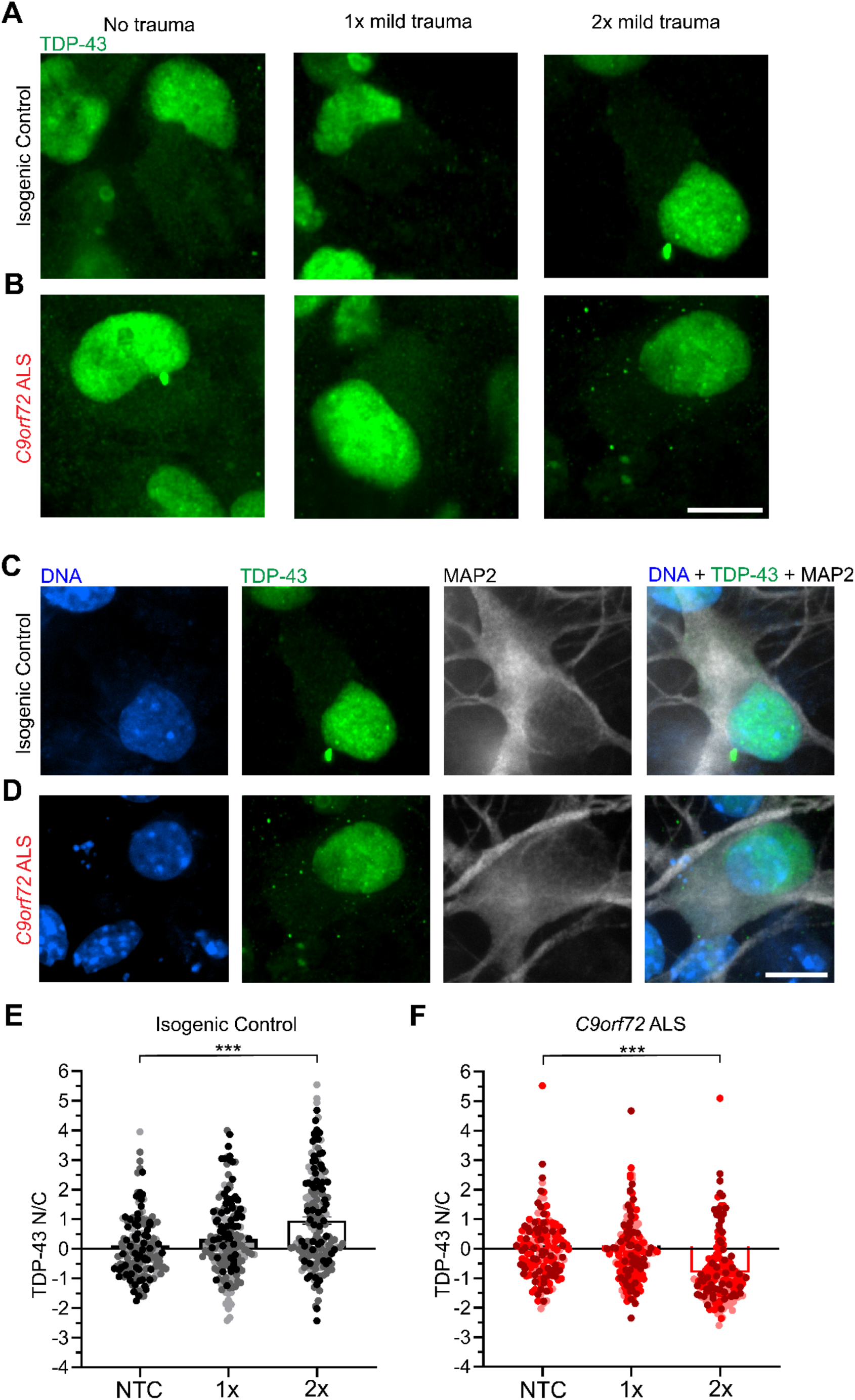
Single channel characterization and individual data point analysis of motor neurons assessing TDP-43 distribution following repetitive mild trauma. (A-B) Visualization of TDP-43 distribution (GFP) without heatmap filter in isogenic control (**A**) or *C9orf72* ALS **(B)** motor neurons. Scale bar = 10 μm. **(C-D)** Single channel image visualization of representative images for isogenic control (**C**) or *C9orf72* ALS (**D**) motor neurons. Scale bar = 10 μm. **(E-F)** Individual data points for experimental averages used in Figure 7 in either isogenic control (**E**) or *C9orf72* ALS (**F**). Data are motor neurons from 3 indpeendent differentiations. Different shades represent different experiments. n = 168-179. Two-way ANOVA for **E** and **F** p < 0.0001. Tukey’s multiple comparison test relative to NTC is indicated. *** = p < 0.005.

## SUPPLEMENTARY VIDEOS

**Supplementary Video 1. High magnification visualization of stretch trauma.** Increased resolution emphasizing vertical and horizontal stretch dynamics of the post array.

**Supplementary Video 2: Induced stretch trauma of a flexible, PDMS-bottomed plate.** Video shows 30FPS high-speed camera capture of a spraypainted wells. The left well is a no-trauma control, while the right well depicts a 2mm displacement. The second video depicts a higher resolution video of the stretched well.

## REFERENCES

1. Brown, R.H. and A. Al-Chalabi, Amyotrophic Lateral Sclerosis. New England Journal of Medicine, 2017. 377(2): p. 162–172.

2. Hardiman, O., et al., Amyotrophic lateral sclerosis. Nature Reviews Disease Primers, 2017. 3: p. 17071–17071.

3. Shaw, P.J., Molecular and cellular pathways of neurodegeneration in motor neurone disease, in Journal of Neurology, Neurosurgery and Psychiatry. 2005, BMJ Publishing Group. p. 1046–1057.

4. Grad, L.I., et al., Clinical Spectrum of Amyotrophic Lateral Sclerosis (ALS), in Cold Spring Harbor perspectives in medicine. 2017.

5. Masrori, P. and P. Van Damme, Amyotrophic lateral sclerosis: a clinical review, in European Journal of Neurology. 2020, Blackwell Publishing Ltd. p. 1918–1929.

6. Zou, Z.Y., et al., *Genetic epidemiology of amyotrophic lateral sclerosis: A systematic review and meta-analysis.* Journal of Neurology, Neurosurgery and Psychiatry, 2017. 88(7): p. 540–549.

7. DeJesus-Hernandez, M., et al., Expanded GGGGCC hexanucleotide repeat in noncoding region of C9ORF72 causes chromosome 9p-linked FTD and ALS. Neuron, 2011. 72(2): p. 245–56.

8. Majounie, E., et al., Frequency of the C9orf72 hexanucleotide repeat expansion in patients with amyotrophic lateral sclerosis and frontotemporal dementia: a cross-sectional study. The Lancet Neurology, 2012. 11.

9. Renton, A.E., et al., A hexanucleotide repeat expansion in C9ORF72 is the cause of chromosome 9p21-linked ALS-FTD. Neuron, 2011. 72(2): p. 257–68.

10. Simón-Sánchez, J., et al., The clinical and pathological phenotype of C9ORF72 hexanucleotide repeat expansions. Brain, 2012. 135(3): p. 723–735.

11. Gendron, T.F., et al., Antisense transcripts of the expanded C9ORF72 hexanucleotide repeat form nuclear RNA foci and undergo repeat-associated non-ATG translation in c9FTD/ALS. Acta Neuropathologica, 2013. 126(6): p. 829–844.

12. Freibaum, B.D. and J.P. Taylor, The role of dipeptide repeats in C9ORF72-related ALS-FTD, in Frontiers in Molecular Neuroscience. 2017, Frontiers Media S.A.

13. Gomez-Deza, J., et al., Dipeptide repeat protein inclusions are rare in the spinal cord and almost absent from motor neurons in C9ORF72 mutant amyotrophic lateral sclerosis and are unlikely to cause their degeneration. Acta neuropathologica communications, 2015. 3: p. 38–38.

14. Lee, Y.B., et al., Hexanucleotide repeats in ALS/FTD form length-dependent RNA Foci, sequester RNA binding proteins, and are neurotoxic. Cell Reports, 2013. 5(5): p. 1178–1186.

15. McEachin, Z.T., et al., RNA-mediated toxicity in C9orf72 ALS and FTD, in Neurobiology of Disease. 2020, Academic Press Inc.

16. Woollacott, I.O.C. and S. Mead, The C9ORF72 expansion mutation: Gene structure, phenotypic and diagnostic issues, in Acta Neuropathologica. 2014. p. 319–332.

17. Zu, T., et al., RAN proteins and RNA foci from antisense transcripts in C9ORF72 ALS and frontotemporal dementia. Proceedings of the National Academy of Sciences of the United States of America, 2013. 110(51).

18. Beck, J., et al., Large C9orf72 hexanucleotide repeat expansions are seen in multiple neurodegenerative syndromes and are more frequent than expected in the UK population. American Journal of Human Genetics, 2013. 92(3): p. 345–353.

19. Benussi, L., et al., C9ORF72 hexanucleotide repeat number in frontotemporal lobar degeneration: A genotype-phenotype correlation study. Journal of Alzheimer’s Disease, 2014. 38(4): p. 799–808.

20. Esselin, F., et al., Clinical Phenotype and Inheritance in Patients With C9ORF72 Hexanucleotide Repeat Expansion: Results From a Large French Cohort. Frontiers in Neuroscience, 2020. 14.

21. Murphy, N.A., et al., Age-related penetrance of the C9orf72 repeat expansion. Scientific Reports, 2017. 7(1).

22. Spargo, T.P., et al., Calculating variant penetrance from family history of disease and average family size in population-scale data. Genome Medicine, 2022. 14(1).

23. Shu, L., et al., The Association between C9orf72 Repeats and Risk of Alzheimer’s Disease and Amyotrophic Lateral Sclerosis: A Meta-Analysis. Parkinson’s Disease, 2016. 2016.

24. Kumar, V., G.M. Hasan, and M.I. Hassan, Unraveling the role of RNA mediated toxicity of C9orf72 repeats in C9-FTD/ALS, in Frontiers in Neuroscience. 2017, Frontiers Media S.A.

25. Leko, M.B., et al., Molecular mechanisms of neurodegeneration related to C9orf72 hexanucleotide repeat expansion, in Behavioural Neurology. 2019, Hindawi Limited.

26. Nordin, A., et al., Extensive size variability of the GGGGCC expansion in C9orf72 in both neuronal and non-neuronal tissues in 18 patients with ALS or FTD. Human Molecular Genetics, 2014. 24(11): p. 3133–3142.

27. Van Blitterswijk, M., M. Dejesus-Hernandez, and R. Rademakers, How do C9ORF72 repeat expansions cause amyotrophic lateral sclerosis and frontotemporal dementia: Can we learn from other noncoding repeat expansion disorders?, in Current Opinion in Neurology. 2012. p. 689–700.

28. Douglas, A.G.L. and D. Baralle, Reduced penetrance of gene variants causing amyotrophic lateral sclerosis. J Med Genet, 2024. 61(3): p. 294–297.

29. Murphy, N.A., et al., Age-related penetrance of the C9orf72 repeat expansion. Sci Rep, 2017. 7(1): p. 2116.

30. Van Wijk, I.F., et al., Assessment of risk of ALS conferred by the GGGGCC hexanucleotide repeat expansion in C9orf72 among first-degree relatives of patients with ALS carrying the repeat expansion. Amyotroph Lateral Scler Frontotemporal Degener, 2024. 25(1-2): p. 188–196.

31. Chen, H., et al., Head injury and amyotrophic lateral sclerosis. American Journal of Epidemiology, 2007. 166(7): p. 810–816.

32. Franz, C.K., et al., Impact of traumatic brain injury on amyotrophic lateral sclerosis: from bedside to bench. Journal of Neurophysiology, 2019. 122(3): p. 1174–1185.

33. Liu, G., et al., *Head Injury and Amyotrophic Lateral Sclerosis: A Meta-Analysis*, in *Neuroepidemiology*. 2021, S. Karger AG. p. 11–19.

34. Burton, D. and M. Aisen, Traumatic Brain Injury, in Handbook of Secondary Dementias. 2006, CRC Press. p. 83–118.

35. Ng, S.Y. and A.Y.W. Lee, Traumatic Brain Injuries: Pathophysiology and Potential Therapeutic Targets, in Frontiers in Cellular Neuroscience. 2019, Frontiers Media S.A.

36. Bramlett, H.M. and W.D. Dietrich, Long-Term Consequences of Traumatic Brain Injury: Current Status of Potential Mechanisms of Injury and Neurological Outcomes. Journal of Neurotrauma, 2015. 32(23): p. 1834–1848.

37. Krishnamoorthy, V. and M.S. Vavilala, Traumatic Brain Injury and Chronic Implications beyond the Brain, in JAMA Network Open. 2022, American Medical Association. p. E229486–E229486.

38. McKee, A.C., et al., The spectrum of disease in chronic traumatic encephalopathy. Brain, 2013. 136(1): p. 43–64.

39. Daneshvar, D.H., et al., *Post-traumatic neurodegeneration and chronic traumatic encephalopathy*, in Molecular and Cellular Neuroscience. 2015, Academic Press Inc. p. 81–90.

40. Rademakers, R., et al., TDP-43 and FUS in amyotrophic lateral sclerosis and frontotemporal dementia, in Lancet Neurol. 2010. p. 995–1007.

41. Scotter, E.L., H.J. Chen, and C.E. Shaw, TDP-43 Proteinopathy and ALS: Insights into Disease Mechanisms and Therapeutic Targets, in Neurotherapeutics. 2015, Springer New York LLC. p. 352–363.

42. Kawakami, I., T. Arai, and M. Hasegawa, The basis of clinicopathological heterogeneity in TDP-43 proteinopathy, in Acta Neuropathologica. 2019, Springer Verlag. p. 751–770.

43. Suk, T.R. and M.W.C. Rousseaux, *The role of TDP-43 mislocalization in amyotrophic lateral sclerosis*, in Molecular Neurodegeneration. 2020, BioMed Central Ltd.

44. Kwong, L.K., et al., *TDP*-43 proteinopathy: The neuropathology underlying major forms of sporadic and familial frontotemporal lobar degeneration and motor neuron disease, in Acta Neuropathologica. 2007. p. 63–70.

45. Liscic, R.M., et al., ALS and FTLD: Two faces of TDP-43 proteinopathy, in European Journal of Neurology. 2008. p. 772–780.

46. De Boer, E.M.J., et al., TDP-43 proteinopathies: A new wave of neurodegenerative diseases, in Journal of Neurology, Neurosurgery and Psychiatry. 2021, BMJ Publishing Group. p. 86–95.

47. McKee, A.C., et al., TDP-43 Proteinopathy and Motor Neuron Disease in Chronic Traumatic Encephalopathy. Journal of Neuropathology and Experimental Neurology, 2010. 69(9): p. 918–929.

48. McKee, A.C. and M.E. Robinson, Military-related traumatic brain injury and neurodegeneration. Alzheimer’s and Dementia, 2014. 10(3 SUPPL.).

49. Tan, X.L., et al., Transactive Response DNA-Binding Protein 43 Abnormalities after Traumatic Brain Injury. Journal of Neurotrauma, 2019. 36: p. 87–99.

50. Sherman, S.A., et al., Stretch Injury of Human Induced Pluripotent Stem Cell Derived Neurons in a 96 Well Format. Scientific Reports, 2016. 6.

51. Finan, J.D., et al., Regional mechanical properties of human brain tissue for computational models of traumatic brain injury. Acta Biomaterialia, 2017. 55: p. 333–339.

52. Phillips, J.K., et al., Method for High Speed Stretch Injury of Human Induced Pluripotent Stem Cell-derived Neurons in a 96-well Format. Journal of Visualized Experiments, 2018. 134.

53. Adams, J.H.G. D; Gennarelli, T; Maxwell, W, *Diffuse axonal injury in non-missile head injury.* Journal of Neurology, Neurosurgery and Psychiatry, 1991. 54(6): p. 481–483.

54. Meythaler, J.M., et al., Current concepts: diffuse axonal injury-associated traumatic brain injury. Arch Phys Med Rehabil, 2001. 82(10): p. 1461–71.

55. Czeiter, E., et al., Traumatic axonal injury in the spinal cord evoked by traumatic brain injury. J Neurotrauma, 2008. 25(3): p. 205–13.

56. Siedler, D.G., et al., Diffuse axonal injury in brain trauma: insights from alterations in neurofilaments. Front Cell Neurosci, 2014. 8: p. 429.

57. del Mar, N., et al., A novel closed-body model of spinal cord injury caused by high-pressure air blasts produces extensive axonal injury and motor impairments. Exp Neurol, 2015. 271: p. 53–71.

58. Sherman, S.A., et al., Stretch Injury of Human Induced Pluripotent Stem Cell Derived Neurons in a 96 Well Format. Sci Rep, 2016. 6: p. 34097.

59. Ziller, M.J., et al., Dissecting the Functional Consequences of De Novo DNA Methylation Dynamics in Human Motor Neuron Differentiation and Physiology. Cell Stem Cell, 2018. 22(4): p. 559–574.e9.

60. Gabler, L.F., J.R. Crandall, and M.B. Panzer, Assessment of Kinematic Brain Injury Metrics for Predicting Strain Responses in Diverse Automotive Impact Conditions. Ann Biomed Eng, 2016. 44(12): p. 3705–3718.

61. Sanchez, E.J., et al., *A reanalysis of football impact reconstructions for head kinematics and finite element modeling.* Clin Biomech (Bristol, Avon), 2019. 64: p. 82–89.

62. Takhounts, E.G., et al., Investigation of traumatic brain injuries using the next generation of simulated injury monitor (SIMon) finite element head model. Stapp Car Crash J, 2008. 52: p. 1–31.

63. Zhao, W., et al., Performance Evaluation of a Pre-computed Brain Response Atlas in Dummy Head Impacts. (1573-9686 (Electronic)).

64. Wu, T., et al., Integrating Human and Nonhuman Primate Data to Estimate Human Tolerances for Traumatic Brain Injury. LID - 071003 [pii] LID - 10.1115/1.4053209 *[doi].* (1528–8951 (Electronic)).

65. Mitevska, A., et al., Polyurethane Culture Substrates Enable Long-Term Neuron Monoculture in a Human in vitro Model of Neurotrauma. Neurotrauma Rep, 2023. 4(1): p. 682–692.

66. Jiang, J., et al., Gain of Toxicity from ALS/FTD-Linked Repeat Expansions in C9ORF72 Is Alleviated by Antisense Oligonucleotides Targeting GGGGCC-Containing RNAs. Neuron, 2016. 90(3): p. 535–550.

67. Westergard, T., et al., Repeat-associated non-AUG translation in C9orf72-ALS/FTD is driven by neuronal excitation and stress. EMBO Mol Med, 2019. 11(2).

68. Anderson, E.N., et al., Traumatic injury induces stress granule formation and enhances motor dysfunctions in ALS/FTD models. Human Molecular Genetics, 2018. 27(8): p. 1366–1381.

69. Anderson, E.N., et al., Traumatic injury compromises nucleocytoplasmic transport and leads to TDP-43 pathology. eLife, 2021. 10.

70. Kiskinis, E., et al., All-Optical Electrophysiology for High-Throughput Functional Characterization of a Human iPSC-Derived Motor Neuron Model of ALS. Stem Cell Reports, 2018. 10(6): p. 1991–2004.

71. Kiskinis, E., et al., Pathways Disrupted in Human ALS Motor Neurons Identified through Genetic Correction of Mutant SOD1. Cell Stem Cell, 2014.

72. Tsioras, K., et al., Analysis of proteome-wide degradation dynamics in ALS SOD1 iPSC-derived patient neurons reveals disrupted VCP homeostasis. Cell Rep, 2023: p. 113160.

73. Farrawell, N.E., et al., Distinct partitioning of ALS associated TDP-43, FUS and SOD1 mutants into cellular inclusions. Scientific Reports, 2015. 5.

74. Mackenzie, I.R.A., et al., Pathological TDP-43 distinguishes sporadic amyotrophic lateral sclerosis from amyotrophic lateral sclerosis with SOD1 mutations. Annals of Neurology, 2007. 61(5): p. 427–434.

75. Melamed, Z.e., et al., Premature polyadenylation-mediated loss of stathmin-2 is a hallmark of TDP-43-dependent neurodegeneration. Nature Neuroscience, 2019. 22(2): p. 180–190.

76. Krus, K.L., et al., Loss of Stathmin-2, a hallmark of TDP-43-associated ALS, causes motor neuropathy. Cell Reports, 2022. 39(13).

77. Klim, J.R., et al., ALS-implicated protein TDP-43 sustains levels of STMN2, a mediator of motor neuron growth and repair. Nat Neurosci, 2019. 22(2): p. 167–179.

78. Prudencio, M., et al., Truncated stathmin-2 is a marker of TDP-43 pathology in frontotemporal dementia. J Clin Invest, 2020. 130(11): p. 6080–6092.

79. Al-Chalabi, A. and O. Hardiman, The epidemiology of ALS: A conspiracy of genes, environment and time, in Nature Reviews Neurology. 2013, Nature Publishing Group. p. 617–628.

80. Oskarsson, B., D.K. Horton, and H. Mitsumoto, *Potential Environmental Factors in Amyotrophic Lateral Sclerosis*, in Neurologic Clinics. 2015, W.B. Saunders. p. 877–888.

81. Bozzoni, V., et al., Amyotrophic lateral sclerosis and environmental factors. Functional Neurology, 2016. 31(1): p. 7–19.

82. Vasta, R., et al., Unraveling the complex interplay between genes, environment, and climate in ALS. The Lancet, 2022. 75.

83. Anderson, E.N., et al., Traumatic injury induces stress granule formation and enhances motor dysfunctions in ALS/FTD models. Hum Mol Genet, 2018. 27(8): p. 1366–1381.

84. Bjorklund, G.R., et al., Traumatic brain injury induces TDP-43 mislocalization and neurodegenerative effects in tissue distal to the primary injury site in a non-transgenic mouse. Acta Neuropathol Commun, 2023. 11(1): p. 137.

85. Evans, T.M., et al., The effect of mild traumatic brain injury on peripheral nervous system pathology in wild-type mice and the G93A mutant mouse model of motor neuron disease. Neuroscience, 2015. 298: p. 410–23.

86. Thomsen, G.M., et al., A model of recurrent concussion that leads to long-term motor deficits, CTE-like tauopathy and exacerbation of an ALS phenotype. J Trauma Acute Care Surg, 2016. 81(6): p. 1070–1079.

87. Thomsen, G.M., et al., Acute Traumatic Brain Injury Does Not Exacerbate Amyotrophic Lateral Sclerosis in the SOD1 (G93A) Rat Model. eNeuro, 2015. 2(3).

88. Wiesner, D., et al., Reversible induction of TDP-43 granules in cortical neurons after traumatic injury. Exp Neurol, 2018. 299(Pt A): p. 15–25.

89. Freibaum, B.D., et al., GGGGCC repeat expansion in C9orf72 compromises nucleocytoplasmic transport. Nature, 2015. 525(7567): p. 129–133.

90. Zhang, K., et al., The C9orf72 repeat expansion disrupts nucleocytoplasmic transport. Nature, 2015. 525(7567): p. 56–61.

91. Barmada, S.J., et al., Cytoplasmic mislocalization of TDP-43 is toxic to neurons and enhanced by a mutation associated with familial amyotrophic lateral sclerosis. J Neurosci, 2010. 30(2): p. 639–49.

92. Zhang, P., et al., Chronic optogenetic induction of stress granules is cytotoxic and reveals the evolution of ALS-FTD pathology. Elife, 2019. 8.

93. Sahana, T.G., et al., *c-Jun N-Terminal Kinase Promotes Stress Granule Assembly and Neurodegeneration in C9orf72-Mediated ALS and FTD*. J Neurosci, 2023. 43(17): p. 3186–3197.

94. Kassouf, T., et al., Targeting the NEDP! enzyme to ameliorate ALS phenotypes through stress granule disassembly. Science Advances, 2023. 9(13).

95. Alam, A., et al., Modeling the Inflammatory Response of Traumatic Brain Injury Using Human Induced Pluripotent Stem Cell Derived Microglia. J Neurotrauma, 2023. 40(19-20): p. 2164–2173.

96. Chaves, R.S., et al., Amyloidogenic Processing of Amyloid Precursor Protein Drives Stretch-Induced Disruption of Axonal Transport in hiPSC-Derived Neurons. J Neurosci, 2021. 41(49): p. 10034–10053.

97. Ramirez, S., et al., Modeling Traumatic Brain Injury in Human Cerebral Organoids. Cells, 2021. 10(10).

98. Snapper, D.M., et al., Development of a novel bioengineered 3D brain-like tissue for studying primary blast-induced traumatic brain injury. J Neurosci Res, 2023. 101(1): p. 3–19.

99. Lai, J.D., et al., KCNJ2 inhibition mitigates mechanical injury in a human brain organoid model of traumatic brain injury. Cell Stem Cell, 2024. 31(4): p. 519–536.e8.

100. Balendra, R., et al., Transcriptome-wide RNA binding analysis of C9orf72 poly(PR) dipeptides. Life Sci Alliance, 2023. 6(9).

101. Parameswaran, J., et al., Antisense, but not sense, repeat expanded RNAs activate PKR/eIF2α-dependent ISR in C9ORF72 FTD/ALS. Elife, 2023. 12.

102. Besnard-Guerin, C., Cytoplasmic localization of amyotrophic lateral sclerosis-related TDP-43 proteins modulates stress granule formation. Eur J Neurosci, 2020. 52(8): p. 3995–4008.

103. Ding, Q., et al., TDP-43 Mutation Affects Stress Granule Dynamics in Differentiated NSC-34 Motoneuron-Like Cells. Front Cell Dev Biol, 2021. 9: p. 611601.

104. Fang, M.Y., et al., Small-Molecule Modulation of TDP-43 Recruitment to Stress Granules Prevents Persistent TDP-43 Accumulation in ALS/FTD. Neuron, 2019. 103(5): p. 802–819 e11.

105. Swarup, V., et al., Abnormal regenerative responses and impaired axonal outgrowth after nerve crush in TDP-43 transgenic mouse models of amyotrophic lateral sclerosis. J Neurosci, 2012. 32(50): p. 18186–95.

106. Jankovic, T., et al., Differential Expression Patterns of TDP-43 in Single Moderate versus Repetitive Mild Traumatic Brain Injury in Mice. Int J Mol Sci, 2021. 22(22).

107. Chou, C.C., et al., TDP-43 pathology disrupts nuclear pore complexes and nucleocytoplasmic transport in ALS/FTD. Nature Neuroscience, 2018. 21(2): p. 228–239.

108. Wangler, L.M. and J.P. Godbout, Microglia moonlighting after traumatic brain injury: aging and interferons influence chronic microglia reactivity. Trends Neurosci, 2023.

109. Ichida, J.K. and E. Kiskinis, Probing disorders of the nervous system using reprogramming approaches. Embo j, 2015.

110. Andersen, J., et al., Generation of Functional Human 3D Cortico-Motor Assembloids. Cell, 2020. 183(7): p. 1913–1929.e26.

111. Velasco, S., et al., Individual brain organoids reproducibly form cell diversity of the human cerebral cortex. Nature, 2019. 570(7762): p. 523–527.

112. Rosenfeld, J.V., et al., Blast-related traumatic brain injury. Lancet Neurol, 2013. 12(9): p. 882–893.

113. Xiong, Y., A. Mahmood, and M. Chopp, Animal models of traumatic brain injury. Nat Rev Neurosci, 2013. 14(2): p. 128–42.

114. Schimmel, S.J., S. Acosta, and D. Lozano, Neuroinflammation in traumatic brain injury: A chronic response to an acute injury. Brain Circ, 2017. 3(3): p. 135–142.

115. Jha, R.M., P.M. Kochanek, and J.M. Simard, Pathophysiology and treatment of cerebral edema in traumatic brain injury. Neuropharmacology, 2019. 145(Pt B): p. 230–246.

116. Franz, C.K., et al., Impact of traumatic brain injury on amyotrophic lateral sclerosis: from bedside to bench. J Neurophysiol, 2019. 122(3): p. 1174–1185.

117. Ortega, J.A., et al., Nucleocytoplasmic Proteomic Analysis Uncovers eRF1 and Nonsense-Mediated Decay as Modifiers of ALS/FTD C9orf72 Toxicity. Neuron, 2020. 106(1): p. 90–107.e13.

